# Cell layer-specific expression of the homeotic MADS-box transcription factor PhDEF contributes to modular petunia petal morphogenesis

**DOI:** 10.1101/2021.04.03.438311

**Authors:** M. Chopy, Q. Cavallini-Speisser, P. Chambrier, P. Morel, J. Just, V. Hugouvieux, Bento S. Rodrigues, C. Zubieta, M. Vandenbussche, M. Monniaux

**Affiliations:** Laboratoire de Reproduction et Développement des Plantes, Université de Lyon, ENS de Lyon, UCB Lyon 1, CNRS, INRAE, 69007 Lyon, France; Laboratoire de Physiologie Cellulaire et Végétale, Université Grenoble-Alpes, CNRS, CEA, INRAE, IRIG-DBSCI, 38000 Grenoble, France

## Abstract

Floral homeotic MADS-box transcription factors ensure the correct morphogenesis of floral organs, which are organized in different cell layers deriving from the meristematic L1, L2 and L3 layers. How cells from these distinct layers acquire their respective identity and coordinate their growth to ensure normal floral organ morphogenesis is unresolved. Here, we study petunia petals that form a limb and tube through congenital fusion, a complex morphology that coevolved with pollinators. We have identified petunia mutants expressing the B-class MADS-box gene *PhDEF* in the epidermis or in the mesophyll of the petal only, called wico and star respectively. Strikingly, wico flowers form a strongly reduced tube while their limbs are almost normal, while star flowers form a normal tube but very reduced and unpigmented limbs, showing that petunia petal morphogenesis is highly modular. Comparative transcriptome analysis of star, wico and wild-type petals revealed a strong down-regulation of the anthocyanin production pathway in star petals including its major regulator *ANTHOCYANIN2 (AN2).* We found that PhDEF directly binds to *AN2* regulatory sequence *in vitro* by gel shift assay, and *in vivo* by chromatin immunoprecipitation, suggesting that PhDEF directly activates the petal pigmentation pathway by activating *AN2*. Altogether, we show that cell-layer specific homeotic activity in petunia petals differently impacts tube and limb development, revealing the relative importance of the different cell layers in the modular architecture of petunia petals.

## INTRODUCTION

All plant aerial organs derive from clonally-distinct layers, named L1, L2 and L3 in the shoot apical meristem (SAM) (Satina et al., 1940). Within the L1 and L2 layers, cells divide anticlinally, thereby maintaining a clear layered structure in all aerial organs produced by the SAM (Meyerowitz, 1997; Stewart and Burk, 1970; Scheres, 2001). Already at the embryonic stage, meristematic cell layers express different genes and have distinct identities (Abe et al., 1999; Lu et al., 1996), that are maintained in the adult SAM (Yadav et al., 2014). During flower development, floral organ identity will be appended on top of layer identity by the combinatorial expression of homeotic floral genes, most of which are MADS-box genes (Coen and Meyerowitz, 1991; Schwarz-Sommer et al., 1990). How these master floral regulators specify all floral organ features, such as organ size, shape, pigmentation, and cellular properties, while maintaining layer-specific identities, is unknown.

Petals are often the most conspicuous organs of the flower, and they display a tremendous diversity in size, shape and pigmentation across flowering plants (Moyroud and Glover, 2017). Floral organ identity is specified by a combination of A-, B-and C-class identity genes as proposed by the classical ABC model established on *Arabidopsis thalian*a (Arabidopsis) and *Antirrhinum majus* (snapdragon), and B-class genes are particularly important for petal identity (Coen and Meyerowitz, 1991; Schwarz-Sommer et al., 1990; Morel et al., 2017). B-class proteins, belonging to MADS-box transcription factors, are grouped in the DEF/AP3 and the GLO/PI subfamilies, named after the snapdragon/Arabidopsis B-class proteins DEFICIENS/APETALA3 and GLOBOSA/PISTILLATA (Purugganan et al., 1995; Theißen et al., 1996). These proteins act as obligate heterodimers consisting of one DEF/AP3 and one GLO/PI protein, and this complex activates its own expression for maintenance of high expression levels all along petal and stamen development (Tröbner et al., 1992). In petunia, gene duplication has generated four B-class genes, namely *PhDEF* and *PhTM6* belonging to the *DEF/AP3* subfamily, and *PhGLO1* and *PhGLO2* belonging to the *GLO/PI* subfamily (Vandenbussche et al., 2004; Rijpkema et al., 2006; van der Krol et al., 1993; Angenent et al., 1992). Mutating the two members of each subfamily (*phdef phtm6* or *phglo1 phglo2* double mutants) results in a classical B-function mutant phenotype with homeotic transformation of petals into sepals and stamens into carpels (Vandenbussche et al., 2004; Rijpkema et al., 2006). Additionally, gene copies within the *DEF/AP3* subfamily have diverged in function: while *PhDEF* exhibits a classical B-class expression pattern largely restricted to developing petals and stamens, *PhTM6* is atypically expressed in stamens and carpels, and its upregulation depends on the petunia C-function genes (Rijpkema et al., 2006; Heijmans et al., 2012a). As a consequence, the single *phdef* mutant displays a homeotic conversion of petals into sepals, while the stamens are normal due to functional redundancy with *PhTM6* (Rijpkema et al., 2006). The petunia *phdef* mutant is therefore an interesting model to study the mechanism of petal identity specification alone since it displays a single-whorl complete homeotic transformation, which is quite rare for floral homeotic mutants that generally show defects in two adjacent whorls.

Flowers from the *Petunia* genus develop five petals, that arise as individual primordia and fuse congenitally (Vandenbussche et al., 2009). Mature petals are fully fused and the corolla is organized in two distinct domains: the tube and the limb. Variation in the relative size of the tube and the limb are observed among wild species of *Petunia*, where flowers with a long tube grant nectar access to long-tongued hawkmoths or hummingbirds, while wide and short tubes are easily accessible to bees (Galliot et al., 2006). The short-and long-tube species cluster separately on a phylogeny of wild *Petunia* species, and the short-tube phenotype is likely the ancestral one (Reck-Kortmann et al., 2014). Pollinator preference assays and field observations have confirmed that tube length and limb size are discriminated by pollinators and thereby might play a role in reproductive isolation, together with multiple other traits of the pollination syndromes such as limb pigmentation or volatile emission (Venail et al., 2010; Hoballah et al., 2007; Galliot et al., 2006). Tube and limb therefore appear to act as different functional modules in the petunia flower.

Although the petunia petal tube and limb seem to play important ecological roles, the mechanisms driving their development are mostly unknown. Tube and limb develop as relatively independent entities in flowers from the Solanaceae family, to which petunia belongs: for instance, tube length and limb width are uncorrelated traits in intra-specific crosses performed in *Nicotiana* or *Jaltomata* (Bissell and Diggle, 2008; Kostyun et al., 2019). Moreover, tube and limb identities can be acquired independently: this is strikingly observed in the petunia *blind* mutant, a partial A-class mutant that forms an almost wild-type tube topped by functional anthers, due to ectopic C-class activity in the second whorl (Cartolano et al., 2007). Apart from the petal identity genes, the molecular players involved in petunia tube or limb morphogenesis are mostly unknown. General growth factors affect petal development as a whole (both tube and limb) together with other vegetative or reproductive traits (Vandenbussche et al., 2009; Terry et al., 2019; Brandoli et al., 2020), but very few genes have been found to specifically affect growth of one subdomain of the petal (Zenoni et al., 2004). Therefore, the mechanisms of petunia tube and limb morphogenesis remain to be fully explored. In contrast, the genetic and molecular bases of petunia petal pigmentation are extremely well characterized, thanks to the plethora of mutants that have been isolated over decades of breeding and research (Bombarely et al., 2016; Tornielli et al., 2009). Petunia limb pigmentation is mainly due to the accumulation of anthocyanins in the vacuole of adaxial epidermal cells. Briefly, the earliest steps of anthocyanin production are ensured by a MBW regulatory complex composed of an R2R3-<u>M</u>YB transcription factor (either ANTHOCYANIN2 (AN2), AN4, DEEP PURPLE (DPL) or PURPLE HAZE), a <u>b</u>HLH transcription factor (AN1 or JAF13), and a <u>W</u>D-40 repeat protein (AN11), which drives the expression of anthocyanin biosynthesis enzymes and proteins involved in vacuolar acidification of epidermal cells (Albert et al., 2011; de Vetten et al., 1997; Spelt et al., 2000; Quattrocchio et al., 1998, 1999, 1993). How this pathway is activated, after regulators such as PhDEF have specified petal identity, has not been elucidated so far.

In this work, we present petunia flowers with strongly affected tube or limb development, that we respectively named wico and star, and that spontaneously arose from plants mutant for *PhDEF*. We provide genetic and molecular evidence that both of these flower types are periclinal chimeras, resulting from the layer-specific excision of the transposon inserted into the *PhDEF* gene, restoring *PhDEF* activity either in the epidermis or in the mesophyll of the petal. The star and wico phenotypes indicate that in the petunia petal, the epidermis mainly drives limb morphogenesis while the mesophyll mainly drives tube morphogenesis. This is seemingly different from previous studies in snapdragon flowers, another species with fused petals, where *def* periclinal chimeras indicated that epidermal *DEF* expression was making a major contribution to overall petal morphology (Perbal et al., 1996; Vincent et al., 2003; Efremova et al., 2001). We characterized in detail the star and wico petal phenotypes at the tissue and cellular scale, and found evidence for non-cell-autonomous effects affecting cell identity between layers. We sequenced the total petal transcriptome from wild-type (wt), wico and star flowers at three developmental stages, and we found that a large proportion of the genes involved in anthocyanin production were downregulated in star petal samples, as could be expected from their white petals. We further showed, by gel shift assay and chromatin immunoprecipitation, that PhDEF binds to the terminator region of *AN2*, thereby likely directly activating its expression and triggering the first steps of limb pigmentation. Our results and our unique flower material promise to improve our understanding of tube and limb morphogenesis in petunia, and address the broader question of how organ identity and cell layer identity superimpose during organ development.

## Results

### Spontaneous appearance of two phenotypically distinct classes of partial revertants from the *phdef-151* locus

Previously described null alleles for the *PhDEF* gene (also named *GP* or *pMADS1*) were obtained by either ethyl methanesulfonate (EMS) mutagenesis (de Vlaming et al., 1984; Rijpkema et al., 2006) or by γ-radiation (van der Krol et al., 1993). Because neither of these alleles were straightforward to genotype in a heterozygous state, we screened our sequence-indexed *dTph1* transposon mutant population in the W138 genetic background (Vandenbussche et al., 2008) for new insertions into *PhDEF*. We identified a new mutant allele named *phdef-151*, referring to the *dTph1* insertion position 151 bp downstream of the ATG in the first exon of the *PhDEF* gene, predicted to fully disrupt the MADS-domain in the protein sequence by premature termination of the first exon due to multiple STOP codons in the different reading frames of *dTph1*. As observed for previously identified *phdef* null alleles, *phdef-151* flowers display a complete homeotic conversion of petals into sepals, while heterozygous or homozygous wild-type siblings display red-coloured wt petals. *phdef-151* is thus very likely a null mutant allele.

While growing homozygous *phdef-151* individuals during several seasons, we repeatedly observed the spontaneous appearance of inflorescence side branches that developed flowers with a partial restoration of petal development (Figure 1, Supplemental Figure 1), suggesting excision of the *dTph1* transposon from the *phdef-151* allele specifically in these sidebranches. Remarkably, these partially revertant flowers could be classified as belonging to either one of two contrasting phenotypic classes, that we named star and wico, and that could even occur simultaneously in different branches on the same plant (Fig. 1A). For both phenotypic classes, we obtained more than 15 independent reversion events. The star flowers (Fig. 1D-F), named in reference to their star-shaped petals, grow an elongated tube similar to wt flowers, but their limbs are underdeveloped: they appear to mainly grow around the mid-vein with strongly reduced lateral expansion, hence losing the typical round shape of wt limb. Moreover, they have almost white petals, suggesting strongly reduced accumulation of anthocyanins. We quantified these changes in flower morphology (Fig. 1K-N) and found that total limb area was reduced almost 5-fold in star flowers (Fig. 1M). In contrast, total tube length was only slightly reduced in star as compared to wt (Fig. 1L), and this was mainly due to a reduction in length of domain D1, corresponding to the part of the tube fused with stamens (as defined in (Stuurman et al., 2004), Fig. 1K), while length of the rest of the tube (domain D2) remained unchanged (Fig. 1L, Supplemental Figure 2). As a result, the ratio between limb area and tube length, which we use as a simple measure for overall corolla morphology, is reduced about 4-fold in star flowers as compared to wt (Fig. 1N). In addition, we occasionally observed fully pigmented secondary revertant sectors of various sizes in the star genetic background, in some cases leading to the development of a single wt-like petal in a star flower background (Fig. 1J). These revertant sectors, observed multiple times, always exhibited simltaneous restauration of pigmentation and normal petal limb growth patterns, demonstrating that the strongly reduced pigmentation in star petals was due to impaired PhDEF function, and not to an additional mutation in the pigmentation pathway.

**Figure 1.**
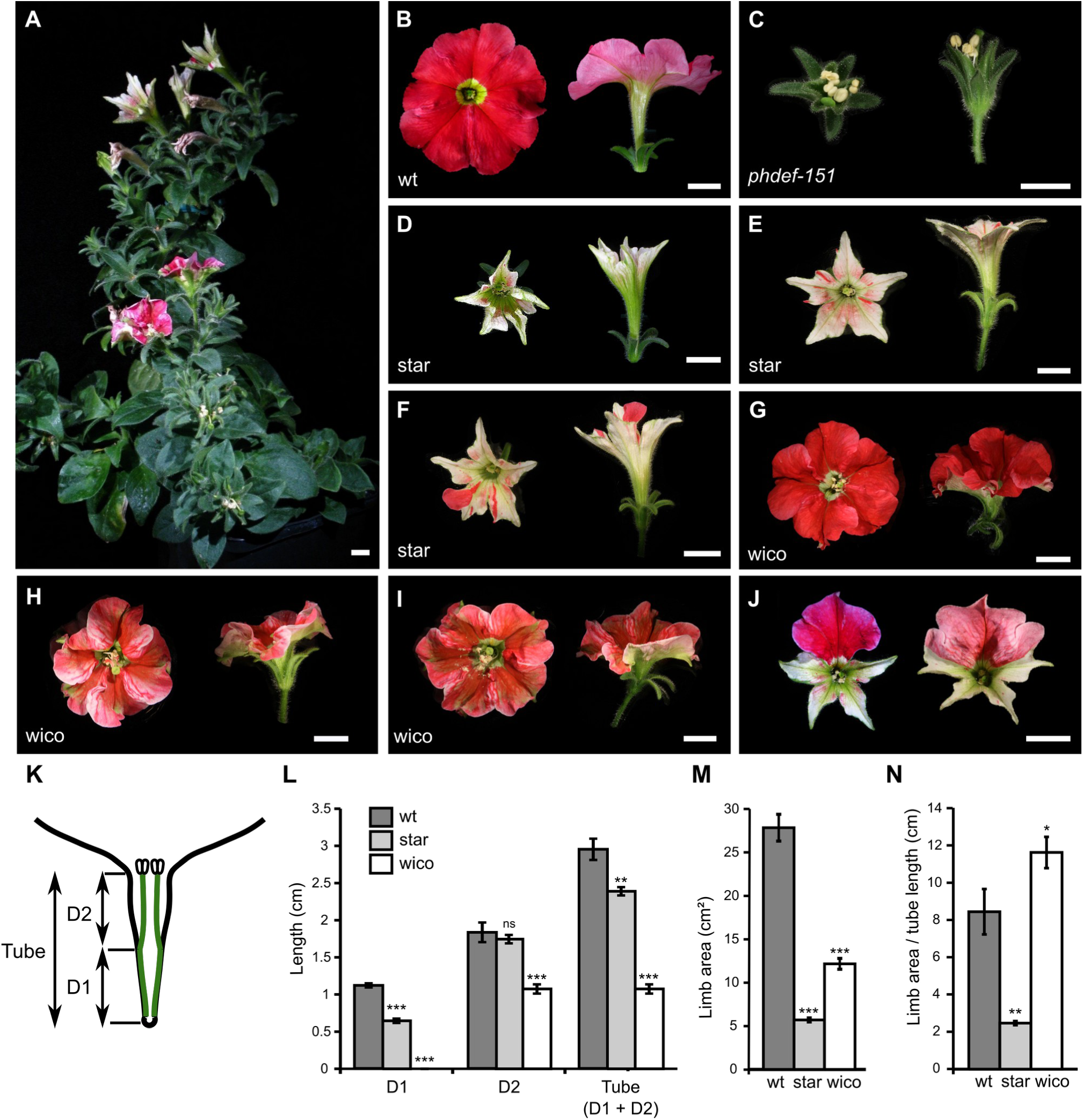
Macroscopic description of the star and wico flowers. **(A)** *phdef-151* mutant plant harboring one branch with wico revertant flowers and one branch with star revertant flowers. Scale bar: 1 cm. **(B-I)** Representative wt (B), *phdef-151* (C), star (D-F) and wico (G-I) flowers from a top (left) and side (right) view. The star and wico flowers come from independent reversion events (from different *phdef-151* plants or from different branches of a single *phdef-151* plant). Scale bar: 1 cm. **(J)** Two star flowers with additional L1-revertant sectors in one petal (left) or one petal and two half petals (right). **(K)** Schematic cross-section of a wt flower, showing stamens (in green) partially fused to the petal tube. The region of the tube fused to stamens is named D1, and the region of the tube where stamens are free is named D2, as defined in (Stuurman et al., 2004). **(L)** Average length of regions D1, D2 and total tube length in wt, star and wico flowers. **(M)** Average limb area in wt, star and wico flowers. **(N)** Average ratio between limb area and tube length in wt, star and wico flowers. n = 7 wt flowers, n = 12 star flowers from 4 different branches, n = 18 wico flowers from 5 different branches. Student’s t test (* p < 0.05, ** p < 0.01, *** p < 0.005). Error bars represent ± s.e.m.

The wico flowers, named after their <u>wi</u>de <u>co</u>rolla, grow round-shaped and pigmented limb while their tube remains underdeveloped (Fig. 1G-I). Limb pigmentation ranged from pink to bright red, and green sepaloid tissue was observed around the mid-veins, commonly well visible in all wico flowers on the abaxial side of the petals (see for instance Supplemental Figure 1E). Total tube length was reduced about 3-fold in wico flowers, with domain D1 being absent since stamens were totally unfused to the tube (Supplemental Figure 2), while domain D2 was significantly reduced in size (Fig. 1L). Limb area was also about 2-fold reduced in wico as compared to wt flowers (Fig. 1M), but the ratio between limb area and tube length was higher than in wt flowers (Fig. 1N), indicating the larger contribution of limb tissue to total corolla morphology in wico flowers. In summary, the star flowers form an almost normal tube but small, misshaped and unpigmented limb, while the wico flowers form almost normally shaped and pigmented limb but a tube strongly reduced in length. These contrasting phenotypes suggest that tube and limb development can be uncoupled in petunia flowers, at least to some degree.

### The star and wico flowers result from excision of the *dTph1* transposon from the *phdef-151* **locus**

Reversion of a mutant phenotype towards a partial or a complete wt phenotype is classically observed in unstable transposon insertion mutant alleles. In the petunia W138 line from which *phdef-151* originates, the *dTph1* transposon is actively transposing (Gerats et al., 1990). We assumed therefore that the star and wico flowers were caused by the excision of *dTph1* from the *PhDEF* locus. *dTph1* transposition is generally accompanied by an 8-bp duplication of the target site upon insertion, and excision can have various outcomes depending on the length and nature of the remaining footprint (van Houwelingen et al., 1999). Hence, we first hypothesized that the distinct star and wico phenotypes were caused by different types of alterations of the *PhDEF* coding sequence after the excision of *dTph1*.

To test this hypothesis, we characterized the *phdef-151* locus from in total 14 star and 14 wico independent reversion events (Figure 2). For this, we extracted genomic DNA from sepals or petals of star and wico flowers, and we amplified the part of the *PhDEF* locus containing the *dTph1* transposon with primers flanking the insertion site (Fig. 2A). All samples produced a mixture of *PhDEF* fragments, some containing the *dTph1* transposon and some where *dTph1* had been excised (Fig. 2B). We specifically sequenced the small fragments resulting from *dTph1* excision in star and wico petal samples, including *phdef-151* second whorl organs as a control (Fig. 2C). In *phdef-151,* the *dTph1*-excised alleles were always out-of-frame, with either 7 or 8 additional nucleotides as compared to the wt sequence. Due to a reading frame shift, both of these alleles are expected to produce an early truncated protein likely not functional (Fig. 2C), in line with the normal *phdef* mutant phenotype observed in these plants. In contrast, in both star and wico flowers we could find either wt sequences (found 1 time and 3 times independently in star and wico flowers respectively) or in-frame footprint alleles consisting of various additions of 6 nucleotides (alleles further named *PhDEF+6*, found 13 times and 11 times independently in star and wico flowers respectively, Fig. 2C). These last insertions are predicted to result in proteins with 2 additional amino-acids inserted towards the end of the DNA-binding MADS domain (Fig. 2C). Together these results demonstrate that wico and star revertant flowers depend on the presence of an in-frame *def-151* derived excision allele that partially restores petal development. However, and in contrast to our initial expectations, there was no correlation between the sequence of the locus after excision and the phenotype of the flower, and both star and wico flowers could be found with a wt *PhDEF* excision allele or with an identical *PhDEF+6* allele (e.g. the 6-bp <u>GTCTGG</u> footprint allele was frequently found both in wico and star flowers). This indicates that the phenotypic difference between the star and wico flowers cannot be explained by a differently modified *PhDEF* sequence after *dTph1* excision. Secondly, since the *phdef* mutation is fully recessive (Vandenbussche et al., 2004), the presence of one transposon mutant allele combined with the wt revertant sequence, normally should lead to wt flowers. Together this implied that another molecular mechanism was causing the difference between wico and star flowers.

**Figure 2.**
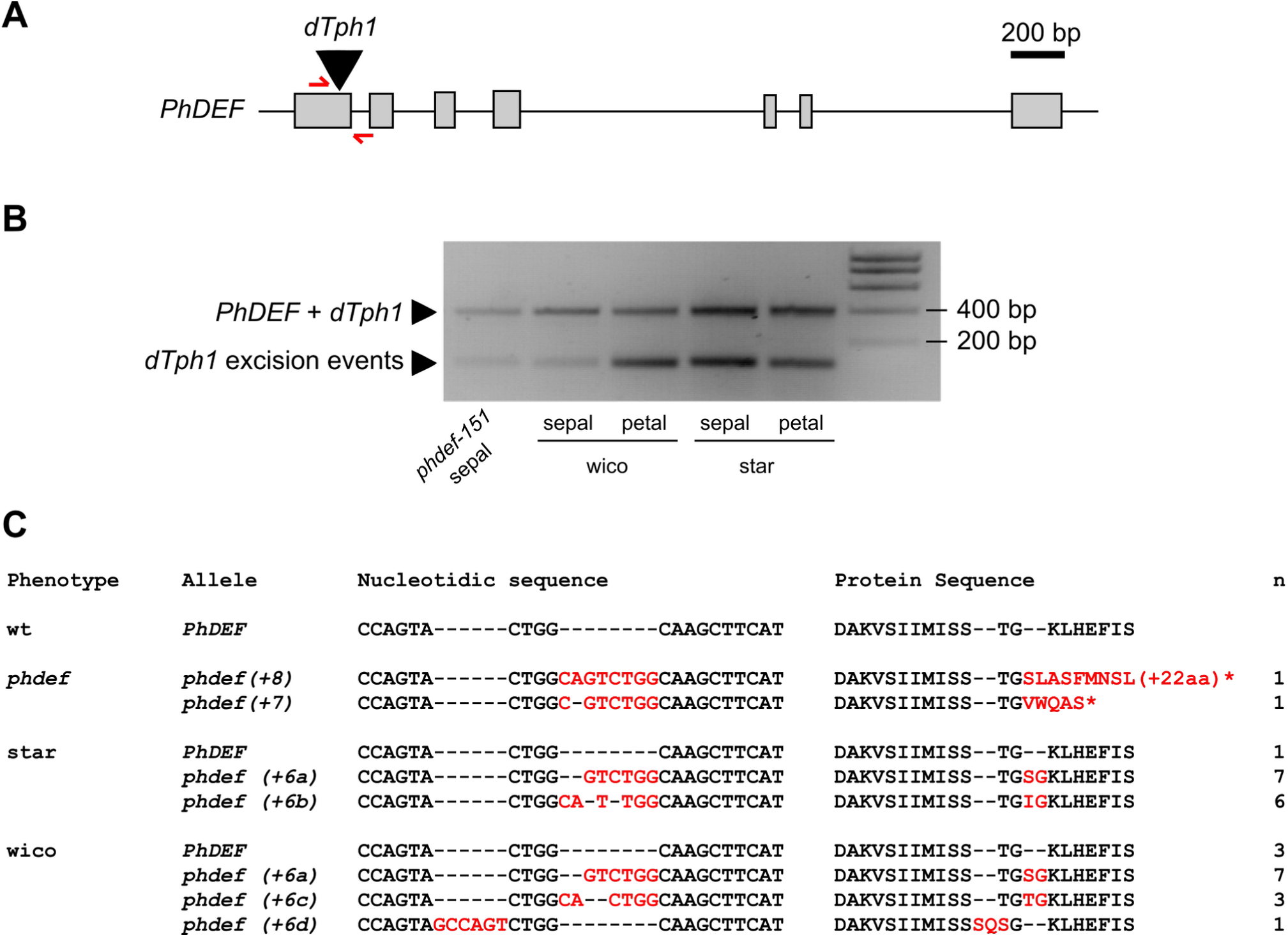
Sequencing the *PhDEF* excision alleles in star and wico flowers. **(A)** *PhDEF* gene model indicating the position of the *dTph1* insertion in the first exon (black triangle) and the primers used for subsequent amplification and sequencing (in red). **(B)** Amplicons generated with primers spanning the *dTph1* insertion site, on genomic DNA from *phdef-151* second whorl organs and star and wico sepals and petals. The large fragment still contains the *dTph1* transposon inserted (expected size: 407 bp), while small fragments result from different events of *dTph1* excision (expected size: 115 bp) and were subsequently sequenced. **(C)** The small *PhDEF* fragments from (B) were sequenced in the second whorl organs of flowers with a *phdef* (n = 2), star (n = 14) and wico (n = 14) phenotype. The nucleotidic sequence and predicted protein sequence are indicated, with STOP codons represented by a star. Additional nucleotides or amino-acids as compared to the wt sequences are indicated in red. n = number of independent reversion events where the same excision footprint was found.

### The wico flowers are L1 periclinal chimeras

Excision of *dTph1* from a gene can occur at different times during plant development: if happening at the zygotic stage, then the whole plant will have a *dTph1*-excised allele. If excision occurs later, this will result in a genetic mosaic (chimera) with a subset of cells carrying the *dTph1* insertion at the homozygous state and others having a *dTph1*-excised allele. This typically leads to branches or flowers with a wt phenotype on a mutant mother plant (supposing a recessive mutation). Furthermore, since all plant organs are organized in clonally-independent cell layers, excision can happen in one cell layer only, thereby creating a periclinal chimera, *i.e.* a branch or flower where cell layers have different genotypes (Frank and Chitwood, 2016; De Keukeleire et al., 2001).

Analyzing the progeny of wico flowers suggested that they were periclinal chimeras, since the wico phenotype turned out not to be heritable (in consequence, they had to be maintained by cuttings of revertant branches). Instead, we found that the progeny of the wico flowers displayed a *phdef* mutant phenotype at a proportion close to 100%, undistinguishable from the parental *phdef-151* allele (Table 1). This suggested that the gametes generated by the wico flowers exclusively carried the mutant *phdef-151* allele, hence resulting in homozygous *phdef-151* mutants in the progeny. Gametes are exclusively derived from the L2 layer in flowering plants (Tilney-Bassett, 1986), therefore indicating that L2-derived germ cells were homozygous mutant for *phdef-151* in wico flowers, which should result in a *phdef* phenotype if the epidermal tissue had the same genotype. This discrepancy suggested that the L1 layer of wico flowers was probably carrying a functional *PhDEF* allele.

**Table 1.** Progeny of the star and wico flowers after selfing. 7 wico flowers and 4 star flowers have been selfed and their progeny has been phenotyped and classified into *phdef*, wt or pink wt phenotype. Summing the star progeny for the 4 parents gives 25 *phdef*, 16 wt and 39 pink wt plants, which is not significantly different to a 1:1:2 ratio (chi-square test, p = 0.22). * For wico, we found 4 plants with wt or pink wt flowers in the progeny, and all of them were linked to the presence of a de novo transposon excision from the *PhDEF* locus, restoring either a *PhDEF+6* (in the case of pink wt progeny) or a wild-type *PhDEF* (in the case of the wt progeny) allele.

|  |  | Phenotype of the progeny (% of the total) |  |  |
| --- | --- | --- | --- | --- |
|  |  | *phdef* | wt | pink wt |
| Parent flower | wico-1 | 15 (94%) |  | 1 (6%) * |
|  | wico-2 | 14 (88%) | 1 (6%) * | 1 (6%) * |
|  | wico-3 | 16 (100%) |  |  |
|  | wico-4 | 15 (94%) |  | 1 (6%) * |
|  | wico-5 | 16 (100%) |  |  |
|  | wico-6 | 12 (100%) |  |  |
|  | wico-7 | 12 (100%) |  |  |
|  | star-1 | 11 (46%) | 4 (17%) | 9 (38%) |
|  | star-2 | 4 (25%) | 4 (25%) | 8 (50%) |
|  | star-3 | 7 (29%) | 5 (21%) | 12 (50%) |
|  | star-4 | 3 (19%) | 3 (19%) | 10 (63%) |

To test this hypothesis, we localized the *PhDEF* transcript in wico flowers by *in situ* hybridization (Figure 3, Supplemental Figure 3). In wt flowers, the *PhDEF* transcript was first detected in the stamen initiation domain, then shortly after in incipient stamen and petal primordia (Fig. 3A, B). At all stages observed, *PhDEF* expression appeared quite homogeneous in all cell layers of the organs, with a stronger expression in the distal part of the petal at later stages of development (Fig. 3C, Supplemental Figure 3). In contrast, in wico flowers *PhDEF* expression was restricted to the L1 and epidermis, all throughout petal development (Fig. 3G-I, Supplemental Figure 3). Therefore, we conclude that wico flowers are the result of an early *dTph1* excision event in one cell from the L1 meristematic layer, resulting in a chimeric flower expressing *PhDEF* only in the epidermis (L1-derived cells) of petals. Wico flowers are therefore L1-periclinal chimeras.

**Figure 3.**
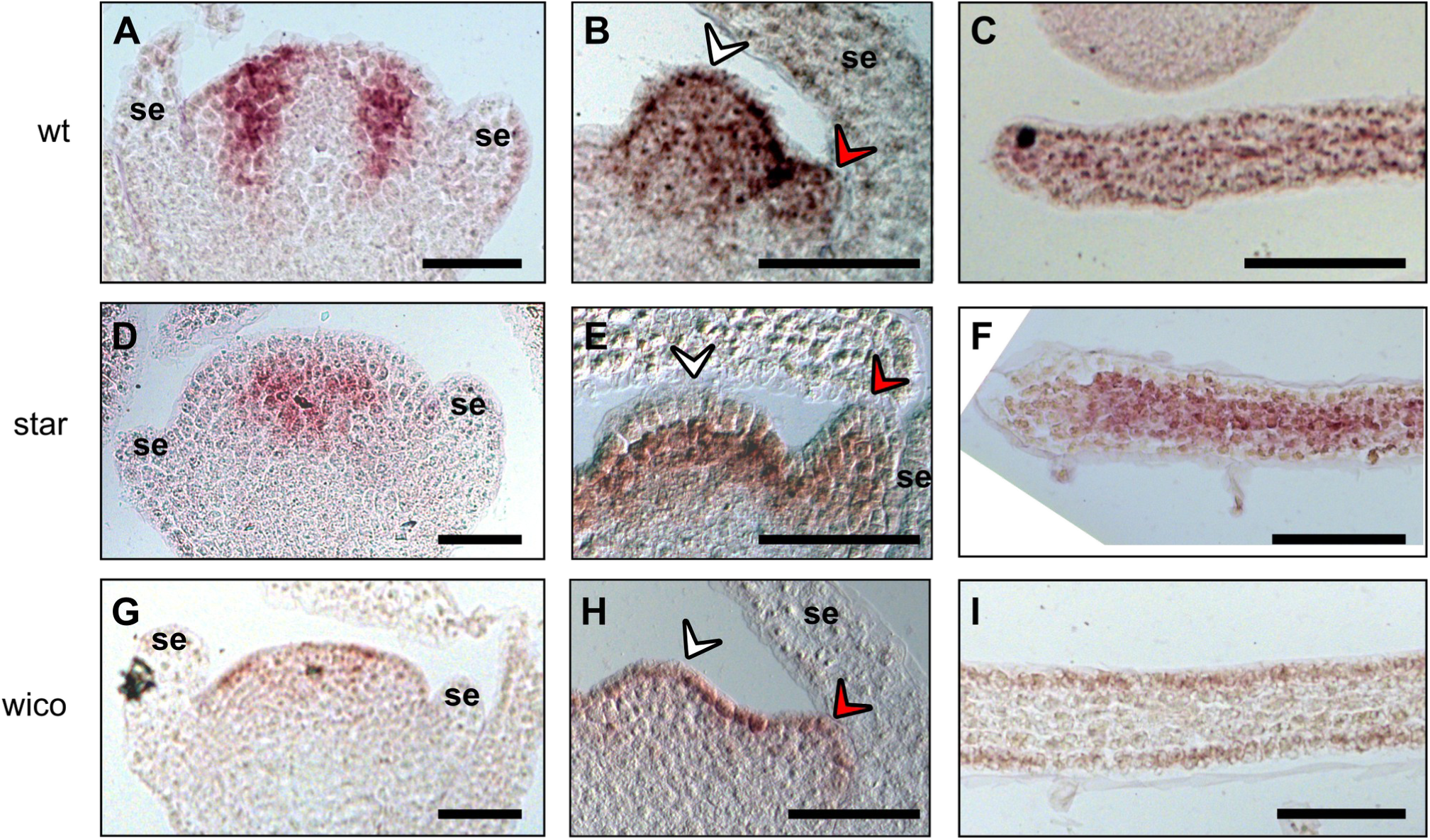
Localization of the *PhDEF* transcript in wt, star and wico flowers by *in situ* hybridization. Longitudinal sections of wt (A, B, C), star (D, E, F) and wico (G, H, I) flowers or young petals hybridized with a DIG-labelled *PhDEF* antisense probe. At the earliest stage chosen (A, D, G), sepals are initiating and *PhDEF* is expressed in the future petal / stamen initiation domain. Note that if the section was not performed at the center of the flower, the *PhDEF* signal might artificially appear to be in the middle of the flower (as in D) whereas it is actually on its flanks. At the middle stage chosen (B, E, H), stamens (white arrowhead) and petals (red arrowhead) are initiating, and *PhDEF* is expressed in both primordia. *PhDEF* expression is also detected at the tip of young petal limb (C, F, I). se: sepals. Scale bar: 50 µm.

### The star flowers are L2 periclinal chimeras

Similarly, we analyzed the progeny of the star flowers, and the star phenotype also turned out not to be heritable, and hence maintained by cuttings of revertant branches. The progeny of the star flowers with a *PhDEF+6* allele yielded three different phenotypic classes (in a proportion close to 1:1:2; Table 1): plants displaying a *phdef* phenotype, plants having wt flowers, and plants carrying flowers with a wild-type architecture but with altered pigmentation, further referred to as « pink wt » (Supplemental Figure 4). We genotyped the *PhDEF* locus in plants descendant from one star parent and carrying flowers with a wt architecture (Supplemental Table 2). We found that all plants with a pink wt phenotype were heterozygous with an out-of-frame *phdef* allele and an in-frame *PhDEF+6* allele, while fully red wt flowers had in-frame *PhDEF+6* alleles at the homozygous state. This indicates that the PhDEF protein with 2 additional amino acids is not 100% fully functional, as it leads to a reduction in limb pigmentation when combined with an out-of-frame allele. The fact that it can ensure normal petal development when at the homozygous state indicates that this is dosage dependent. In summary, the segregation ratio shows that the star gametes carried either the *phdef-151* allele or an in-frame *PhDEF* allele at a 1:1 ratio, and hence that the germ cells generating these gametes were heterozygous for these two alleles. This suggested that in star flowers, the L2 layer was carrying a functional *PhDEF* allele while the L1 layer was homozygous mutant for *phdef-151*.

In support of this, in star flowers *PhDEF* expression was absent from the L1 and epidermis (Fig. 3D-F, Supplemental Figure 3). At the petal margins, underlying layers were also devoid of *PhDEF* expression (Fig. 3F), which likely corresponds to the restricted petal area where cells of L1 origin divide periclinally and invade the mesophyll (Satina and Blakeslee, 1941a). Therefore, we conclude that star flowers are the result of an early *dTph1* excision event in one cell from the L2 meristematic layer, resulting in a chimeric flower expressing *PhDEF* only in the mesophyll (L2-derived cells) of petals. Star flowers are therefore L2-periclinal chimeras. Considering the star and wico phenotypes, we can conclude that the petal epidermis is the main driver for limb morphogenesis (growth, shape and pigmentation), while the mesophyll mainly drives tube morphogenesis (growth and shape).

### Non-cell-autonomous effects of layer-specific *PhDEF* expression on cell identity

Having determined the genetic basis of the star and wico phenotypes, we next wondered how layer-specific *PhDEF* expression affects the determination of cell identity, in the layer where *PhDEF* is expressed (cell-autonomous effect) but also in the layer devoid of *PhDEF* expression (non-cell-autonomous effect). For this, we observed petal adaxial epidermal cells by scanning electron microscopy, and mesophyll cells on petal cross-sections, in wt petals and sepals, and in star and wico petals (Figure 4).

**Figure 4.**
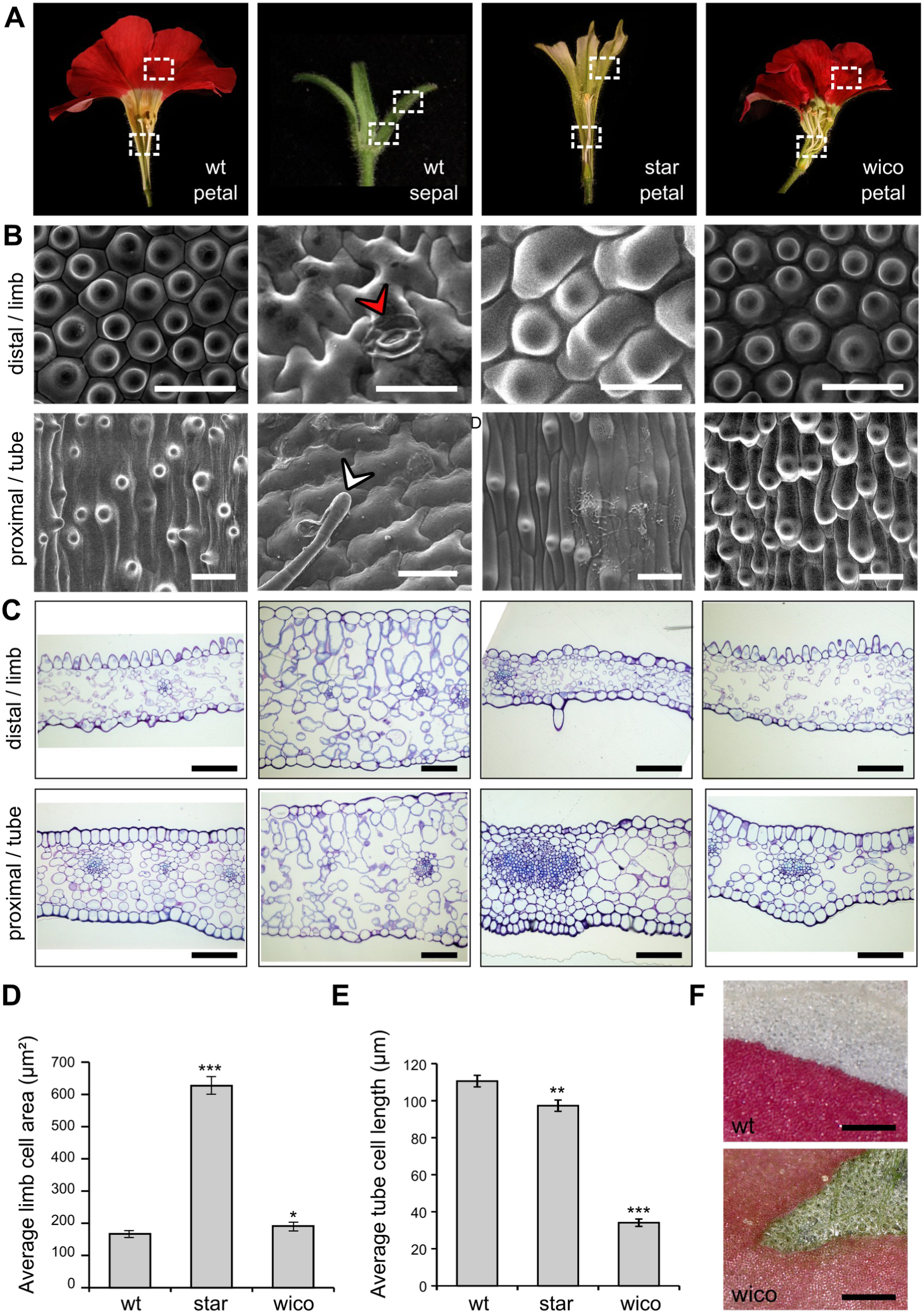
Epidermal and mesophyll cell identities in wt petals and sepals, and star and wico petals. **(A)** From left to right: wt petals, wt sepals, star petals and wico petals cut open longitudinally to show areas used for scanning electron microscopy and cross-sections. Petals were subdivided into limb and tube area, and sepals were subdivided into a distal and a proximal part, as shown by the dotted white rectangles. **(B)** Representative scanning electron micrographs from the adaxial side of a wt petal, wt sepal, star petal and wico petal (from left to right). The red arrowhead points to a stomata and the white arrowhead points to a trichome. Scale bar: 30 µm. **(C)** Representative cross-sections from wt petals, wt sepals, star petals and wico petals (from left to right) stained with toluidine blue. Scale bar: 100 µm. **(D)** Average limb cell area from the adaxial side of wt, star and wico petals (n = 30 cells). Student’s t test (* p < 0.05, ** p < 0.01, *** p < 0.005). **(E)** Average tube cell length from the adaxial side of wt, star and wico petals (n = 45 cells). Wilcoxon rank sum test (* p < 0.05, ** p < 0.01, *** p < 0.005). Error bars represent ± s.e.m. **(F)** Limb area from wt (top) and wico (bottom) petals, after their adaxial epidermis was manually peeled. For wt, the upper half of the picture shows the white underlying mesophyll. For wico, the green triangular area shows the green (chloroplastic) underlying mesophyll.

On the adaxial side of the wt petal, cells from the limb are round and adopt the classical conical shape found in many angiosperm petal limb, while cells from the tube are elongated with a central cone (Fig. 4B) (Cavallini-Speisser et al., 2021). In contrast, the adaxial epidermis of wt sepals (indistinguishable from *phdef-151* second whorl organs) displays typical leaf-like features (Morel et al., 2019), with puzzle-shaped cells interspersed with stomata and trichomes (Fig. 4B). Epidermal cell identity can thus be clearly determined on the basis of cell shape. In wico petals, epidermal limb cells are conical, similar to wt cells from the same area, although marginally bigger (Fig. 4B, D). In contrast, cells from the tube, albeit displaying a similar shape than wt cells, are strongly reduced in length (Fig. 4B, E). This suggests that, in addition to the absence of the D1 region of the tube (Fig. 1K, Supplemental Figure 2), a defect in cell elongation in the D2 region is (at least partly) responsible for overall tube length reduction in wico petals. In star petal tubes, epidermal cells have a similar appearance as in a wt petal tube but are slightly less elongated (Fig. 4B, E). In contrast, epidermal cells from the star limb are very different to both wt petal conical cells and wt sepal puzzle cells: they are slightly bulging cells, more or less roundish, and about 3-times larger than wt conical cells (Fig. 4D). Together with the observation that the star limb area is around 5-times smaller compared to wt limb (Fig. 1M), this indicates that normal activity of PhDEF in the epidermis is required not only for the typical conical cell shape but also for a high cell division rate during petal development. As mentioned earlier, we occasionally observed pigmented revertant sectors on star flowers, resulting from an additional independent *dTph1* excision in the epidermis, generating wt sectors on a star flower (Fig. 1I). These sectors allow the immediate comparison between star and wt epidermal cells on a single sample, confirming the difference in conical cell size, shape and colour (Supplemental Figure 5). Moreover, the star limb adaxial epidermis occasionally forms trichomes (Supplemental Figure 5), a feature that is normally not observed in the wt limb adaxial epidermis. Altogether, these observations suggest that epidermal cells from star limb have an intermediate identity between petal and sepal cells. Since star petals do not express *PhDEF* in their epidermis, these observations show that non-cell-autonomous effects are at stake to specify cell identity.

Mesophyll cell identity was investigated by analyzing petal cross-sections stained with toluidine blue (Fig. 4C). In the wt petal tube, mesophyll cells are big and round and the tissue is loosely arranged. In the limb, mesophyll cells appear smaller and with a more elongated shape, while still very loosely arranged. Sepal mesophyll cells are bigger than petal mesophyll cells, and they display the typical leaf mesophyll organization with an upper palisade layer (elongated and parallel cells) and a lower spongy layer (dispersed cells). Hence mesophyll cell size, shape and tissue-level organization are characteristic features allowing to distinguish between sepal and petal mesophyll tissue. In star petals, we observed no visible difference in mesophyll cell features with wt petals, suggesting that petal cell identity is normally specified in the star petal mesophyll. In wico petals, mesophyll cells also appeared similar to wt: they were round and big in the tube, and slightly more elongated in the limb. Their organization was clearly distinct from the one found in sepals since no palisade layer was observed. However, peeling the epidermis from wico limb revealed that the underlying mesophyll was chloroplastic, similar to a sepal mesophyll and in striking contrast with the white mesophyll of wt petal limb (Fig. 4F). Thus, the *phdef* mutant mesophyll in wico flowers has an intermediate identity between sepal and petal, suggesting again the existence of non-autonomous effects influencing cell identity across layers. The interpretation of these effects is summarized in Supplemental Figure 6.

We wondered if the non-cell-autonomous effects that we observed between layers in the star petals were also happening within a single layer. The revertant sectors observed on star flowers showed a very abrupt transition between pigmented and non pigmented epidermal cells, together with a quite sharp transition in conical cell shape and size (Supplemental Figure 5). In particular, we found a clear file of pigmented cells on a star petal and the scanning electron micrograph revealed that these cells were also conical, in stark contrast with the surrounding flat elongated cells of the star petal mid-vein (Supplemental Figure 5). Therefore, we conclude that within the epidermal layer, cell shape and pigmentation are defined cell-autonomously, suggesting that different processes are at stake for cell-cell communication across and within layers.

### Transcriptome sequencing of star and wico petals

To better understand the molecular basis for the star and wico phenotypes, we performed RNA-Seq on total petal tissue at three developmental stages, including wt and *phdef-151* samples (Figure 5). We chose an early stage (stage 4 as defined in (Reale et al., 2002)), an intermediate stage (stage 8) when tube length is at half its final size, and a late stage (stage 12) before limb is fully expanded (Fig. 5A). For *phdef-151* we only sequenced second-whorl sepal tissue at stage 12 (before anthesis). Principal component analysis showed that developmental stage is the first contributor to variation in gene expression, while genotype corresponds to the second axis of variation (Fig. 5B). All genotypes clustered separately except wico and wt samples which were highly similar at the two later stages. We analyzed one-to-one differential gene expression between mutant and wt samples with DESeq2 (Love et al., 2014) and we found on average 5,818 deregulated genes in *phdef-151*, as compared to 1,854 and 1,115 deregulated genes in star and wico respectively, when averaging for all stages (Fig. 5C, Supplemental Dataset 1). There were generally more upregulated genes than downregulated ones in mutant or chimeric genotypes, and the number of deregulated genes increased with ageing of the petal in both star and wico (Fig. 5C). At stage 12, a large proportion of genes (58-61%) deregulated in wico or star petals were also deregulated in *phdef-151* (Fig. 5D), as expected since wico and star flowers are mutant for *PhDEF* in one cell layer. Genes uniquely deregulated in star or wico flowers represented 36% of deregulated genes for each, and only 16-29% of deregulated genes were jointly deregulated in star and wico flowers, consistent with the very different phenotypes of these flowers. These proportions indicate that the star and wico phenotypes are mostly subtended by the deregulation of sets of genes also deregulated in *phdef-151*, together with the deregulation of a unique set of genes for each genotype.

**Figure 5.**
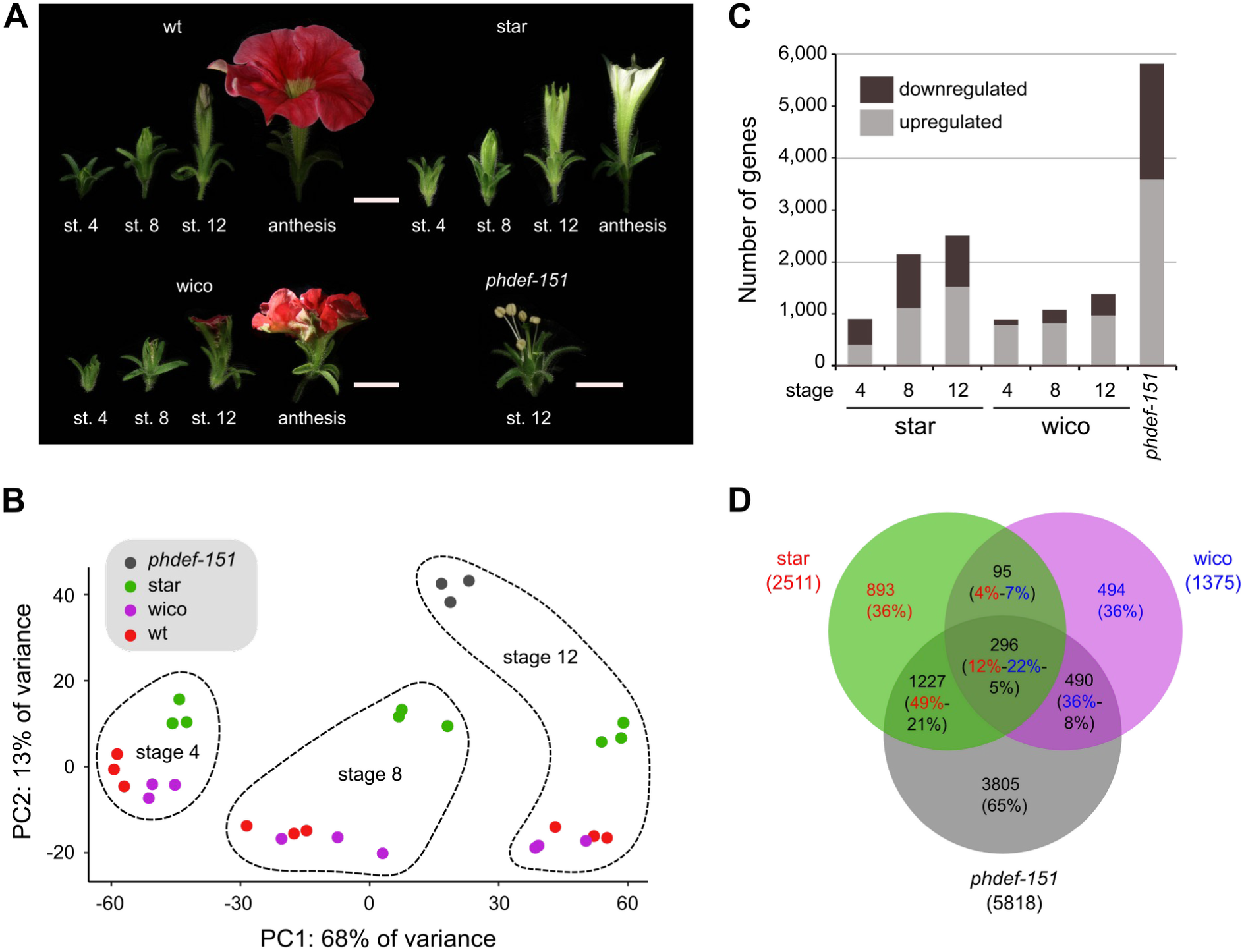
Gene deregulation in star and wico petals. **(A)** Flowers from wt, star, wico and *phdef-151* at stages 4, 8 and 12 (only stage 12 for *phdef-151*), whose petals or sepals were harvested for transcriptome sequencing. Flowers at anthesis are shown for comparison. Scale bar: 1 cm. **(B)** Principal Component Analysis plot of the samples after analysis of variance with DESeq2, showing that the first principal component corresponds to the developmental stage and the second principal component corresponds to the genotype. **(C)** Number of upregulated and downregulated genes in star, wico and *phdef-151*, as compared to wt at the corresponding stages. **(D)** Venn diagram recapitulating the number of deregulated genes in star, wico and *phdef-151* petal samples at stage 12, as compared to wt, and their different intersections. Each sector contains the number of deregulated genes, and between parenthesis is the percentage of genes that it represents from the total number of deregulated genes in the corresponding sample, with a colour code (red = percentage of deregulated genes from star samples / blue = from wico samples / black = from *phdef-151* samples).

In star and wico petals, we found that *PhDEF* was down-regulated about two-fold at all stages (Supplemental Figure 7), as expected since *PhDEF* is expressed in one cell layer only. In contrast, *PhTM6* was not deregulated in star and wico nor in *phdef-151* (Supplemental Figure 7), as expected since this atypical B-class gene is mostly expressed in stamens and carpels and its upregulation depends on the C-function genes (Rijpkema et al., 2006; Heijmans et al., 2012b). Unexpectedly, we observed that the B-class genes *PhGLO1* and *PhGLO2* were not down-regulated in wico petals, and only modestly in star petals, although their expression was almost null in the *phdef-151* mutant (Supplemental Figure 7). The fact that *PhGLO1/PhGLO2* expression does not strictly mirror the expression of *PhDEF* in star and wico petals, which is what we would have expected since the B-class heterodimers are known to activate their own expression, suggests that *PhGLO1/PhGLO2* expression is not entirely dependent on the B-class heterodimeric complexes, in particular in the epidermal layer of the petal.

### PhDEF directly binds *in vivo* to the terminator region of *AN2,* encoding a major regulator of petal pigmentation

The transcriptomes of star and wico petals constitute a promising dataset to identify genes involved in the establishment of petal epidermis and mesophyll identities, and in tube and limb development.

To evaluate their potential to decipher the gene regulatory networks underlying petal development, we decided to focus our attention on genes involved in petal pigmentation. Indeed, the players and regulatory pathways involved in anthocyanin biosynthesis in the petal epidermis have been extremely well described but their relationship with the specifiers of petal identity, to whom PhDEF belongs, is so far unknown. The absence of pigmentation in star petals, the restoration of pigmentation in L1-revertant sectors and the phenotype of the pink wt flowers prompted us to investigate the direct link between *PhDEF* expression and petal pigmentation. For this, we examined the 451 genes down-regulated in both *phdef-151* and star samples (at any stage) but not deregulated in wico samples (Supplemental Dataset 2), and we found 23 anthocyanin-related genes in this gene set (Supplemental Figure 7), out of a total of 42 in the whole genome, which constitutes an exceptionally high enrichment for this gene function (p < 0.001, Fisher’s exact test). We payed a particular attention at the genes possibly involved in the first steps of anthocyanin production, ie encoding proteins involved in the MBW complexes activating anthocyanin biosynthesis (AN1, AN2, AN4, AN11, JAF13, DPL and PURPLE HAZE). We found that *AN1*, *AN2*, *DPL* and *JAF1* were downregulated both in *phdef-151* and star samples (Supplemental Figure 7, Supplemental Dataset 2). DPL is involved in the limb venation pattern (Albert et al., 2011; Zhang et al., 2021) and JAF13 has only a moderate contribution to limb pigmentation (Bombarely et al., 2016), therefore we decided to focus our attention on the two major activators of anthocyanin biosynthesis AN1 and AN2 (Figure 6). Indeed, the *an1* mutant has fully white petals and the *an2* mutant has strongly reduced limb pigmentation (Quattrocchio et al., 1999; Spelt et al., 2000). Furthermore, *AN2* was shown to act as an upstream activator of *AN1* since overexpressing *AN2* in petunia leaves is sufficient to activate *AN1* expression, and for anthocyanins to accumulate (Quattrocchio et al., 1998; Spelt et al., 2000). We observed that both genes were already expressed at stage 4 of wt petal development, before any pigmentation is visible, and their expression levels strongly increased from stage 4 to stage 12, while both being strongly downregulated in star petals and *phdef-151* second whorl organs, but not in wico flowers (Fig. 6A, B). *AN2* was expressed at higher levels than *AN1* at all stages, consistent with its most upstream role in the anthocyanin pigmentation pathway We aimed to test if PhDEF could be a direct activator of *AN1* or *AN2* expression. For this, we first attempted to predict PhDEF binding on the genomic sequences of *AN1* and *AN2*. We used the high-quality transcription factor (TF) binding profile database Jaspar (Fornes et al., 2020; Sandelin et al., 2004), using position weight matrices for each TF to compute relative binding scores that reflect *in vitro* binding preferences (Stormo, 2013). The exact DNA-binding specificity of PhDEF has not been characterized, but the one of its Arabidopsis homologs AP3 and PI has (Riechmann et al., 1996b). However, since PhDEF DNA-binding specificity might be slightly different to those of AP3 and PI, we decided to predict binding for all MADS-box TFs available in Jaspar 2020, accounting for 23 binding profiles including those of AP3 and PI (Fornes et al., 2020). We hypothesized that sequences predicted to be bound by several MADS-box TFs might be high-confidence CArG boxes (the binding site for MADS-box proteins). As a validation of this strategy, we analyzed the genomic sequence of *PhDEF* and found a high-confidence CArG box in the *PhDEF* promoter (visible by the presence of good predicted binding sites for several MADS-box proteins and therefore appearing as a clear black line in Fig. 6C). This CArG box is extremely conserved between distantly-related flowering plants (Rijpkema et al., 2006) and it was shown to be important for *AP3* petal-specific expression and for its auto-activation in Arabidopsis (Hill et al., 1998; Wuest et al., 2012), and for DEF function and binding to its own promoter in Anthirrhinum (Schwarz-Sommer et al., 1992). We next applied this predictive approach to the genomic sequences of *AN1* and *AN2*. For *AN1*, we predicted a high-confidence CarG box (*AN1-bs1*) with a very high score for several MADS-box proteins and for AP3 and PI in particular, in the terminator region (Fig. 6D). For *AN2*, we also predicted one promising candidate binding site (*AN2-bs3*), again in the terminator region of the gene (Fig. 6E), although its binding score was more modest in comparison to *AN1-bs1*.

**Figure 6.**
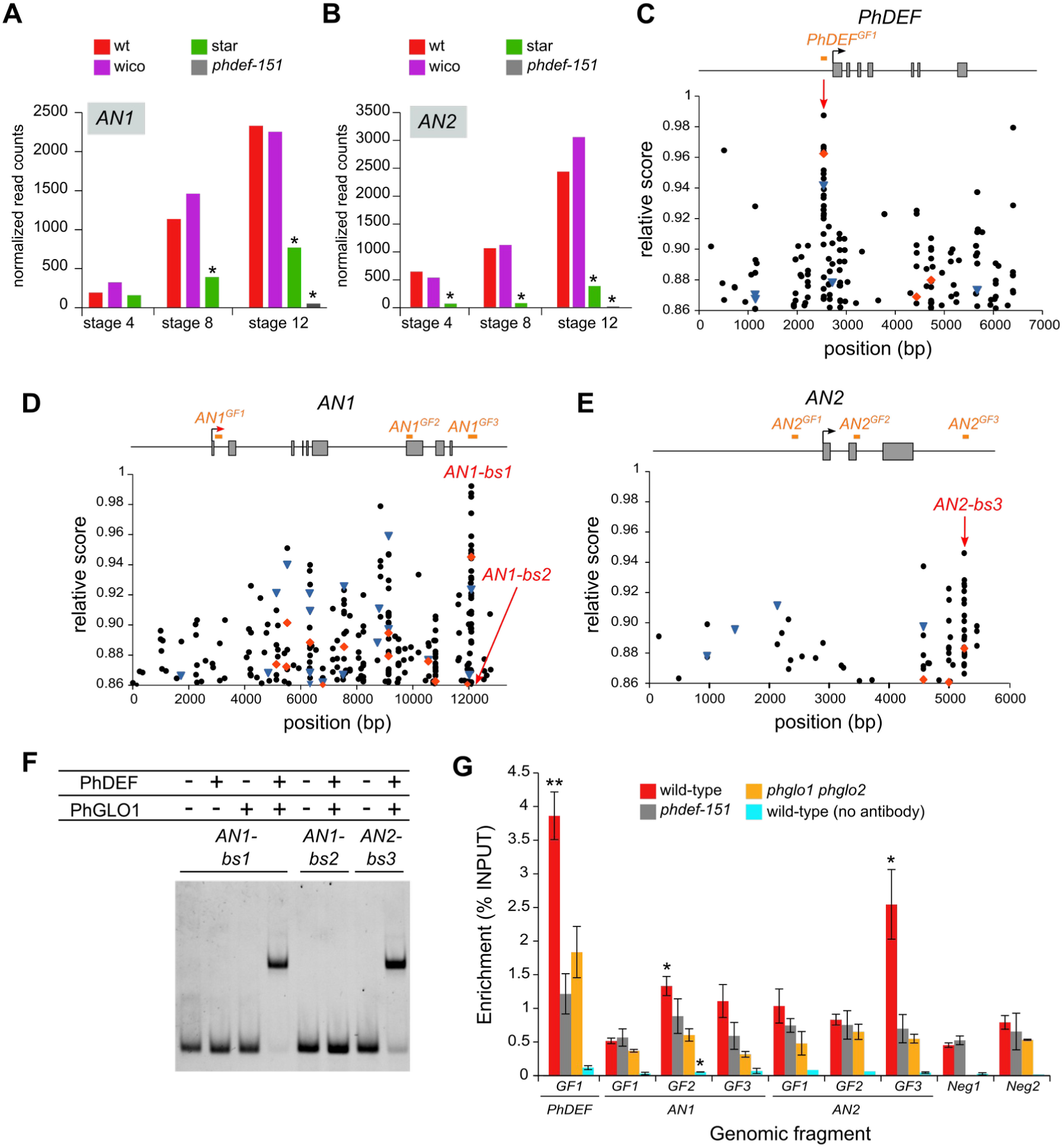
PhDEF binds to *AN2* regulatory region *in vitro* and *in vivo*. **(A, B)** Expression (as normalized read counts calculated by DESeq2) of *AN1* (A) and *AN2* (B) in wt, star, wico and *phdef-151* second whorl organs at stages 4, 8 or 12. Stars indicate significant down-regulation (log2FC < -1 and adjusted p-value < 0.01). **(C-E)** Relative score profiles for AP3 (red diamond), PI (blue triangle) and all other MADS-box TFs (black dots) available on Jaspar, on the genomic sequences of *PhDEF* (C), *AN1* (D) and *AN2* (E). The relative score is computed using the position weight matrix of each TF and is between 0 and 1; only relative scores higher than 0.86 are shown here. The gene model is represented above the score profile with exons as grey rectangles, the transcription start site as an arrow, and the gene model is aligned with the position of the predicted binding sites. For *PhDEF*, the position of a high-confidence CArG box, as explained in the main text, is indicated by a red arrow. In red, are indicated the positions of the sites tested by gel shift in (F): putative PhDEF binding sites (*AN1-bs1* and *AN2-bs3*) and a negative control with a low predicted binding score (*AN1-bs2)*. In orange, are depicted the genomic fragments tested by chromatin immunoprecipitation in (G). **(F)** Representative electrophoretic mobility shift assay (EMSA) gel performed with a combination of *in vitro*-translated PhDEF and/or PhGLO1 proteins, and Cy5-labelled *AN1-bs1, AN1-bs2* or *AN2-bs3* DNA fragments, whose position is depicted in (C-E). **(G)** Enrichment (as percentage of INPUT) of binding of PhDEF to different genomic regions of the chromatin purified from wt, *phdef-151* or *phglo1 phglo2* second whorl organs at stage 8, after immunoprecipitation with an anti-PhDEF directed antibody. The control without antibody was performed on chromatin isolated from wt petals. The position of the genomic fragments tested is depicted in (C-E). Neg1 and Neg2 represent two negative control fragments located in the promoter region of genes not deregulated in the *phdef-151* mutant, and present on different chromosomes than *PhDEF*, *AN1* and *AN2.* For unknown reasons, the Neg1 control region could not be amplified in the *phglo1 phglo2* samples. Stars indicate a significant enrichment of test regions over the average of the two negative control regions for each chromatin sample (one-tailed t-test, * p<0.05, ** p<0.005; n = 3 biological replicates for wt and *phdef-151*, 2 biological replicates for *phglo1 phglo2* and the control without antibody).

To determine if PhDEF could indeed bind to *AN1-bs1* and *AN2-bs3* and potentially regulate *AN1* and *AN2* expression, we performed gel shift assays using *in vitro* translated PhDEF and/or PhGLO1 proteins (Fig. 6F). We found that, when incubating a 60-bp fragment containing *AN1-bs*1 in its center with either PhDEF or PhGLO1, no shift in migration was visible, indicating that neither protein could bind to this site alone. However, when incubating *AN1-bs1* with both PhDEF and PhGLO1 proteins, we observed a clear shift in migration, consistent with the obligate heterodimerization of these proteins necessary for DNA binding (Riechmann et al., 1996a). Similarly, a 60-bp fragment containing *AN2-bs3* in its center, and incubated with PhDEF and PhGLO1 proteins, resulted in a clear shift in migration. In contrast, a control 60-bp fragment named *AN1-bs2*, located in the *AN1* terminator region but predicted to have a very low binding score (relative score under 0.8 both for AP3 and PI), was not bound by the PhDEF + PhGLO1 protein complex, showing that our assay was specific. Therefore PhDEF, when dimerized with PhGLO1, is able to bind to sites in putative regulatory regions in *AN1* and *AN2*, suggesting that it might directly regulate the expression of these two genes.

Next, we tested if PhDEF could bind *in vivo* to genomic regions containing *AN1-bs1* and *AN2-bs3* by chromatin immunoprecipitation (ChIP). We produced recombinant PhDEF protein devoid of its highly conserved MADS domain, to avoid cross-reactivity with other MADS-box proteins, and generated a polyclonal antibody against this truncated PhDEF protein. We performed the ChIP assay on second whorl organs (petal or sepal) from wt, *phdef-151* or *phglo1 phglo2* plants at an intermediate stage of development (stage 8). In wt petal samples, we found a significant binding enrichment for some of the genomic fragments (GF) that we tested, and in particular *PhDEF^GF1^*(Fig. 6G), containing the high-confidence CArG box previously described (Fig. 6C), which is expected since PhDEF activates its own expression. We also observed a significant binding enrichment in *AN2^GF3^* (Fig. 6G), containing the previously identified *AN2-bs3* binding site (Fig. 6E). In contrast, no strong enrichment was detected in any of the *AN1* genomic fragments, even the one containing the *AN1-bs1* strong *in vitro* binding site for PhDEF (AN1^GF3^). Our ChIP assay was specific, since no enrichment was detected for the *phdef-151* mutant, nor for the *phglo1 phglo2* mutant (Fig.6G). The *phglo1 phglo2* samples constitute an indirect control for PhDEF binding, since the PhDEF protein partners PhGLO1/PhGLO2 are absent, thereby indirectly preventing PhDEF binding on DNA. The fact that we do not detect any binding enrichment in these plants shows that our ChIP assay is robust. Therefore, we conclude that PhDEF binds to the terminator region of *AN2 in planta*.

## Discussion

In this work, we identified periclinal chimeras expressing the B-class MADS-box gene *PhDEF* in different cell layers of the flower. This layer-specific expression resulted in the correct development of sub-domains of the petal only, showing that epidermal *PhDEF* expression mainly drives limb morphogenesis while its expression in the mesophyll is more important for tube morphogenesis. This indicates that cell layer-specific actions of PhDEF are different and contribute in a complementary fashion to overall petal development.

### Contribution of cell layers to mature petunia petals

The SAM of all flowering plants is organized in three independent layers. Generally, it is assumed that L1-derived cells form the epidermis, L2-derived cells produce the mesophyll and sub-epidermal tissue, and L3-derived cells generate the ground tissues (inner mesophyll, vasculature, pith of the stem). However, there is variation to this general pattern between organs; for instance Arabidopsis sepals, stamens and carpels derive from these three layers, while petals derive from the L1 and L2 layers only (Jenik and Irish, 2000). Moreover, the contribution of cell layers can vary between the same organ in different species: for instance Datura petals are derived from all three layers, in contrast to petals from Arabidopsis (Satina and Blakeslee, 1941b). Finally, even in one organ from a single species, cell layer contribution is not always homogeneous in different parts of the organ: in Datura petals, the L3 only participates to the vasculature at the base of the organ but does not contribute to the distal part of the petal, and the L1 invades the mesophyll at the petal edges (Satina and Blakeslee, 1941b).

In fact, the contribution of cell layers to mature organ organization can only be strictly assessed by clonal analysis, where one follows cell lineage using trackable cell-autonomous markers. In petunia, no clonal analysis has been performed so far, hence one can only assume which cell layers participate to petal development based on clonal analyses performed in closely-related species. In Datura, member of the Solanaceae family like petunia, periclinal chimeras induced by colchicine treatment and refined histological observations have provided a detailed clonal analysis for cell layers in floral organs (Satina and Blakeslee, 1941b). The first visible event of petal initiation is a periclinal cell division from the L2 layer, and further growth of the petal depends primarily on cell divisions from the L2, both anticlinal and periclinal. The L3 layer only contributes to the vascular tissue at the very base of the petal. L1-derived cells form the epidermis by anticlinal divisions, except at the petal edges where periclinal divisions are observed, leading to L1-derived cells invading the mesophyll. Hence, the Datura petal is formed by all 3 layers with a major contribution of the L1 and L2 layers, and a relative enrichment in L1-derived cells (by thinning of the mesophyll) as we progress from the base towards the tip of the petal. In this work, we hypothesized that the petunia petal is formed similarly. Consistently, we only obtained two phenotypic classes of periclinal chimeras, star and wico, suggesting that L3-specific *PhDEF* expression probably only leads to a *phdef* mutant phenotype.

The contribution of L1-and L2-derived tissues is heterogeneous in the petunia petal. Indeed, cross-sections in the middle of the petal tube indicate that the mesophyll is thick, with several layers of cells (Fig. 4C). The mesophyll tissue is quite dense in this part of the tube, with lacunae between cells being relatively small. In contrast in the limb, mesophyll cells are very small and interspersed with large lacunae. There is also a general thinning of the mesophyll as we progress from the base of the petal towards its edges, whereas the epidermis always appears as a single layer of tightly connected cells. Therefore, it is rather logical that in the petal limb, whose mesophyll is extremely reduced, morphogenesis is driven by the epidermal layer. However, one could not have easily guessed that tube morphogenesis would be mostly driven by the petal mesophyll.

### Different cell layers drive tube and limb morphogenesis

The star and wico phenotypes revealed that in petunia petals, the epidermis is the main driver for limb morphogenesis while the mesophyll is the main driver for tube morphogenesis. The epidermis has been proposed to be the layer in control of organ morphogenesis, since it is a layer under tension that restricts growth of the underlying inner tissues that tend to expand (Kutschera and Niklas, 2007). In particular, epidermal expression of the brassinosteroid receptor BRI1 is sufficient to restore normal leaf morphogenesis in a *bri1* mutant (Savaldi-Goldstein et al., 2007). Similarly, the expression of the auxin transporter PIN1 in the L1 of the SAM is sufficient to restore normal phyllotaxis in a *pin1* mutant (Kierzkowski et al., 2013). However, pieces of evidence suggest that organ inner layers can have an active role in morphogenesis: for instance, mesophyll-specific expression of *ANGUSTIFOLIA* (*AN*) is sufficient to restore normal leaf width in the Arabidopsis *an* mutant (Bai et al., 2010); leaf shape is controlled by the L2-and L3-derived tissues in tobacco (McHale and Marcotrigiano, 1998); and the leaf mesophyll is the main player for leaf flatness in Arabidopsis (Zhao et al., 2020). Moreover, expressing *BRI1* in the root phloem also restaures *bri1* plant dwarfism (Graeff et al., 2020). The contribution of cell layers to organ morphogenesis is thus a complex process that varies between organs, species and the genetic systems investigated.

Our work has confirmed that the petunia petal has a modular structure, since tube and limb can develop relatively independently from each other in the star and wico flowers. This modularity is consistent with previous observations in the literature (described in the Introduction), and in line with the different ecological roles of the tube and the limb for the interaction with pollinators. Our results highlight that a homeotic factor, PhDEF, can participate to the establishment of this modular structure. Indeed, although PhDEF is normally present in all cell layers of the wild-type petal, its action in the different cell layers is mainly responsible for tube or limb development. This provides a possible mechanism, at the tissue level, for the establishment of the modular structure of petunia petals by homeotic genes. It also participates to the understanding of how homeotic genes can specify at the same time the overall identity of an organ and the coordinated development of its different functional modules.

One may wonder if our findings apply outside of petunia flowers. In snapdragon and Arabidopsis flowers, periclinal chimeras for orthologs of *PhDEF* (*DEF* and *AP3* respectively) or *PhGLO1/PhGLO2* (*GLO* and *PI* respectively) have been previously obtained (Perbal et al., 1996; Vincent et al., 2003; Efremova et al., 2001; Bouhidel and Irish, 1996; Jenik and Irish, 2001; Urbanus et al., 2010b). In snapdragon, expression of *DEF* only in the L1 layer largely restores petal development, particularly in the limb, in contrast to the L2/L3 specific *DEF* or *GLO* expression which causes reduced limb growth (Perbal et al., 1996; Vincent et al., 2003; Efremova et al., 2001). Petals are fused into a tube in snapdragon flowers, but the tube is much more reduced than in petunia, hence conclusions on tube length restoration in the chimeras were not drawn by the authors. However, in light of our results, it is clear that snapdragon chimeras expressing *DEF* or *GLO* in the L2/L3 layers restore tube development to a higher degree than limb development, similar to what we observed. In Arabidopsis that has simple and unfused petals, petal size was not fully restored when *AP3* was expressed in one cell layer only, while petal shape was normal (Jenik and Irish, 2001; Urbanus et al., 2010b); in contrast epidermal expression of *PI* was sufficient to restore normal petal development (Bouhidel and Irish, 1996). Therefore, it seems that the contribution of different cell layers to petal development varies across species and depending on the petal identity gene under investigation.

### Autonomous and non-autonomous effects of *PhDEF* expression on petal traits

Our study revealed that petal traits were affected differently by layer-specific *PhDEF* expression (Fig. S6). For instance, epidermal pigmentation is a clearly autonomous trait, since star petals are not pigmented except when wt revertant sectors arise. On the contrary, epidermal cell shape appears to behave as a partially autonomous trait since star epidermal cells have an intermediate phenotype between wt petal conical cells and sepal epidermal cells. Finally, organ size and shape are specified non-autonomously in sub-domains of the petal: *PhDEF* expression in the L1 or L2 is sufficient to specify correct shape of the limb or correct size and shape of the tube respectively, suggesting that in these petal domains, layer-specific *PhDEF* expression is sufficient to signal cells from the other layer to grow normally. The mechanisms for this inter-layer communication remain unknown. Our attempts to detect the PhDEF protein in petal tissue by immuno-histochemistry have been unsuccessful, therefore we do not know if the PhDEF protein itself might be moving between layers, which would be the simplest mechanistic explanation for the non-autonomous traits that we observe. Indeed, in Antirrhinum petals expressing *DEF* in the L2/L3 layers, the DEF protein was found in the epidermis and it is likely why petals from these chimeras are pigmented (Perbal et al., 1996), hence suggesting that no such movement occurs in the star petals that are mostly white. In contrast, Arabidopsis AP3 and PI GFP-fusion proteins are unable to move between cell layers, although they can move within the epidermal layer (Urbanus et al., 2010a, 2010b). In any case, even if the PhDEF protein would move between layers in our chimeric flowers, it is likely to be in small amounts only, otherwise both flower types would have a wt phenotype. Therefore, it is unlikely to be the reason for tube and limb correct development in the star and wico flowers. Alternatively, the non-autonomous effects that we observed might be triggered by mechanical signals transmitted between layers. For instance, in star flowers normal growth of the mesophyll could merely drag along epidermal cells, since cells are connected by their cell walls, which could be sufficient to trigger their expansion and division. Other features, like conical cell shape, might be directly influenced by mechanical signals. Indeed, conical cells are shaped by a circumferential microtubule arrangement controlled by the microtubule-severing protein KATANIN, and altering this arrangement affects conical cell shape (Ren et al., 2017). Microtubule arrangement responds to mechanical signals (Hamant et al., 2008), which are likely to be transmitted between layers. Therefore, it is possible that the formation of bulging cells in the star epidermis is merely triggered by mechanical signals from the growing underlying layer, independent of any petal identity specifier, as was recently evidenced from the observation of conical-like bulges on the hypocotyl of the tubulin kinase mutant *nek6* (Takatani et al., 2020). The molecular or physical nature of the signals involved in communication between layers remains to be explored in full depth.

### Towards the gene regulatory networks of petal development

Our star and wico material granted the opportunity to explore the gene regulatory networks driving petal development in petunia, more specifically by decoupling tube vs. limb development on one hand, and epidermis vs. mesophyll development on the other hand. However, these effects are confounded in our dataset, since both epidermis and limb development are affected in star flowers, whereas both mesophyll and tube development are affected in wico flowers. Further analyses, like for instance sequencing the transcriptome from star and wico limb and tube tissues separately, would help uncouple these effects, but it is not obvious to clearly separate these different domains during early stages of development, which are crucial stages for petal morphogenesis. Spatial transcriptomics techniques would be ideal to precisely dissect transcriptional changes between layers and domains of the petal at young developmental stages. Still, we exploited our transcriptomic dataset by focusing our analysis on anthocyanin-related genes, because the molecular link between the early establishment of petal identity by homeotic transcription factors, such as PhDEF, and the late establishment of petal maturation traits, such as anthocyanin accumulation, was unknown. For this, we examined the presence of anthocyanin-related genes among genes downregulated both in star and *phdef-151* samples, but not deregulated in wico samples. We found a very strong enrichment of anthocyanin-related genes in this dataset, suggesting that the initial triggering event for most of the anthocyanin production pathway was missing in star flowers.

Finally, we investigated the direct link between PhDEF and petal pigmentation and found that, *in vitro*, the PhDEF + PhGLO1 protein complex directly binds to good predicted binding sites in the regulatory regions of *AN1* and *AN2*. We confirmed that PhDEF binds to the genomic region of *AN2 in plant*a by ChIP, but binding to *AN1* was not observed, confirming that *in vitro* binding does not necessarily imply *in vivo* binding, the last being strongly influenced by the local chromatin landscape. The binding site of PhDEF on *AN2* lies in the terminator region of the gene (and the next gene on the chromosome is more than 100 kb away), which although unusual, is not incompatible with an activating role in transcription, through DNA looping to the promoter (Jash et al., 2012) or by promoting transcription termination and reinitiation (Wang et al., 2000). Together with the fact that *AN2* expression is strongly down-regulated in the *phdef-151* transcriptome, our data indicates that PhDEF directly activates *AN2* expression in the petal. Ectopic expression of *AN2* in petunia leaves is sufficient to trigger anthocyanin accumulation in this tissue, by inducing *AN1* expression among others (Spelt et al., 2000; Quattrocchio et al., 1998). Therefore, the fact that PhDEF activates *AN2* expression should be sufficient to launch the whole pigmentation pathway in the wt petal limb.

A direct link between petal identity and pigmentation has never been evidenced before, although genetic evidence in orchid flowers strongly implied that different B-class proteins heteromeric complexes are responsible for specific pigmentation spots in the different petal types, but physical binding of these B-class protein complexes on pigmentation genes was not tested (Hsu et al., 2021). The direct target genes of B-class proteins have been identified by ChIP-Seq and transcriptomic analyses in Arabidopsis (Wuest et al., 2012), but this species has unpigmented petals, thereby preventing to draw any possible link between petal identity and pigmentation. Therefore, to our knowledge, our results show for the first time the direct activation of a petal pigmentation regulator by a petal homeotic gene, which contributes to fill the « missing link » between the identity of a floral organ and its final appearance (Dornelas et al., 2011).

## METHODS

### Plant material, growth conditions and plant phenotyping

The *phdef-151* plants were obtained from the *Petunia x hybrida* W138 line and were grown in a culture room in long day conditions (16h light 22°C; 8h night 18°C; 75-WValoya NS12 LED bars; light intensity: 130 µE). The wico and star flowers were repeatedly obtained from several different *phdef-151* individuals and were maintained by cuttings. Plant and flower pictures were obtained with a CANON EOS 450D camera equipped with objectives SIGMA 18-50mm or SIGMA 50mm. To measure tube length, the flower was cut longitudinally and photographed from the side. To measure limb area, the limbs were flattened as much as possible on a glass slide covered with transparent tape and photographed from the top. The photographs were used to measure D1 and D2 lengths and limb area with ImageJ.

### Genotyping

Extraction of genomic DNA from young leaf tissue was performed according to Edwards et al., 1991. The region spanning the *dTph1* insertion site in *PhDEF* was amplified using primers MLY0935/MLY0936 (Table S1). PCR products were separated on a 2% agarose gel, fragments of interest were purified using the NucleoSpin® Gel and PCR Clean-up kit (Macherey-Nagel), and sequenced with Eurofins SupremeRun reactions.

### *In situ* RNA Hybridization

Floral buds from wt, 2 wico and 1 star lines were fixated overnight in FAA (3.7% formaldehyde, 5% acetic acid, 50% ethanol), cleared in Histo-clear and embedded in paraffin to perform 8 µm sections. *PhDEF* cDNA was amplified from wt petunia inflorescence cDNAs with primers MLY1738/MLY1739 (Supplementary Table 1), generating a 507 bp fragment excluding the part encoding the highly conserved DNA-binding domain. The digoxigenin-labeled RNA probe was synthesized from the PCR fragment by *in vitro* transcription, using T7 RNA polymerase (Boehringer Mannheim). RNA transcripts were hydrolyzed partially for 42 min by incubation at 60°C in 0.1 M Na_2_CO_3_/NaHCO_3_ buffer, pH 10.2. Later steps were performed as described by (Cañas et al., 1994). For imaging, slides were mounted in Entellan (Sigma) and imaged with a Zeiss Axio Imager M2 light microscope equipped with a Zeiss Axio Cam HRc camera.

### Petal cross-sections

Small pieces (around 5 mm^2^) of tissue were harvested from the proximal and distal parts of wt mature sepals, and from the tube and limbs of wt, star and wico mature petals. Samples were fixated overnight in FAA (3.7% formaldehyde, 5% acetic acid, 50% ethanol) and dehydrated in an ethanol series. Preinfiltration was performed in a 1:1 mixture of ethanol:Technovit 7100 (Electron Microscopy Sciences) for 4 h under light agitation, then overnight in a 1:3 ethanol:Technovit 7100 mixture. Infiltration was performed in the infiltration solution for 1.5 h under vacuum, then for one night followed by one additional week. Samples were disposed in the moulds with the polymerization solution for 2 h at room temperature, then mounted with the Technovit 3040 resin to relieve the blocks from the moulds. Blocks were sectioned with a microtome to generate 3-7 µm-thick sections. Slides were incubated for 10 minutes in a 0.1% toluidine blue solution and imaged with a Zeiss Axio Imager M2 light microscope equipped with a Zeiss Axio Cam HRc camera.

### Scanning Electron Microscopy (SEM)

Scanning electron micrographs were obtained with a HIROX SH-1500 bench top environmental scanning electron microscope equipped with a cooling stage. Samples were collected and quickly imaged to limit dehydration, at −5°C and 5 kV settings. For cell area and length measurements, pictures were taken from 3 petal tubes and 3 petal limbs from different wt, star and wico flowers. For each sample, 3 pictures were taken and 5 cells (for the tube) or 10 cells (for the limb) were measured for each picture. Measures were performed with ImageJ by manually drawing the outline or length of the cells.

### RNA-Seq

Petal tissue was collected at 1 pm from several plants stemming from a single star line, a single wico line, and several individual wt plants (progeny of a single star flower) and *phdef-151* plants (progeny of the same star flower). Tube length was macroscopically measured to compare stages, the corolla was cut open and stamens were removed as much as possible from the corolla by pulling on the filaments fused to the tube. One biological replicate contains total petal tissue from 2 flowers. Tissue was grounded in liquid nitrogen and RNA was extracted with the Spectrum Plant Total RNA Kit (Sigma) including on-column DNase digestion (Sigma). RNA integrity and quantity were determined by a Bioanalyzer RNA 6000 Nano assay (Agilent). Libraries were prepared with poly-A enrichment and single-end 75-bp sequencing was performed on a NextSeq 500 platform (Illumina). 16 to 23 million reads were recovered per library. Reads were checked for quality with FastQC v0.11.4 (https://www.bioinformatics.babraham.ac.uk/projects/fastqc/), adaptors and low-quality ends were trimmed with Cutadapt v 1.16 (Martin, 2011) and custom Perl scripts. The reference genome sequence used for transcriptome analysis is the *Petunia axillaris* v1.6.2 HiC genome published in (Bombarely et al., 2016) and further scaffolded by HiC by DNAzoo (Dudchenko et al., 2017, 2018); gene annotations were transferred from the published assembly to the HiC-scaffolded version using Blat (Kent, 2002), Exonerate (Slater and Birney, 2005) and custom Perl scripts. In the rare cases when gene annotations from the published genome mapped to several regions in the HiC-scaffolded genome, these different putative genes were identified by a letter added at the end of the gene identifier (for instance Peaxi162Scf00179g00121a). The complete set of reads was mapped on the reference genome sequence using HISAT2 v2.2.1 (Kim et al., 2015) to identify splicing sites, before performing mapping sample per sample. Reads per gene were counted using FeatureCounts v1.5.1 (Liao et al., 2014). DESeq2 version 3.12 (Love et al., 2014) was used with R version 4.0.3 to perform the Principal Component Analysis and the differential gene expression analysis. Genes having less than 10 reads in the sum of all samples were considered as non-expressed and discarded. Genes were considered to be deregulated if log2FoldChange > 1 or < -1, and p-adjusted value < 0.01. Venn diagrams were built with InteractiVenn (Heberle et al., 2015). Due to the automatic gene name annotation pipeline used in (Bombarely et al., 2016) based on homology with tomato proteins, many of the previously characterized petunia genes have not been annotated according to their first described name, making interpretation of some of the RNA-Seq results less straightforward. We have manually added annotations of 42 genes from the anthocyanin biosynthesis pathway based on the Supplementary Note 7 from (Bombarely et al., 2016), and 31 type-II MIKC-C MADS-box genes based on previous studies from the literature; these annotations can be found in the Supplemental Datatset 1 of this manuscript. We noticed that the gene annotations from three major pigmentation genes, *DFR* (Peaxi162Scf00366g00630), *CHSa* (Peaxi162Scf00047g01225) and *PH1* (Peaxi162Scf00569g00024) were lost during the gene annotation transfer procedure, because they lie in regions of the genome that are still badly resolved. Therefore, we manually searched the position of these transcripts in the HISAT2 output and we were able to map part of the *DFR* and *CHSa* genes to two small scaffolds, while *PH1* position was not found. We added the transcript positions of *DFR* and *CHSa* in the gtf/gff files before running FeatureCounts. The read counts for *DFR* and *CHSa* reported in Supplemental Figure 7 are therefore an under-estimation of their actual expression levels, since we miss part of the genes.

### Prediction of MADS-box TF binding sites

Genomic sequences from *AN1*, *AN2* and *PhDEF* from the *Petunia x hybrid*a R27 line, starting 3 kb upstream the START codon and ending 1 kb downstream the STOP codon, were scanned with all MADS-box TF matrices included in the Jaspar 2020 database (http://jaspar.genereg.net), only removing matrices from AGL42 and AGL55 which are much shorter than the other matrices and therefore yield much higher scores. Relative scores above 0.86 were plotted against their genomic position.

### Electrophoretic Gel Shift Assays (EMSAs)

CDS sequences from *PhDEF* and *PhGLO1* were amplified from *Petunia x hybrida* R27 inflorescence cDNAs with primers MLY2382/MLY2383 and MLY2384/2385 respectively (Supplementary Table 1) and cloned into the *in vitro* translation vector pSPUTK (Stratagene) by NcoI/XbaI restriction. From these vectors, the PhDEF and PhGLO1 proteins were produced with the TnT SP6 High-Yield Wheat Germ Protein Expression System (Promega) according to the manufacturer’s instructions. The terminator regions from *AN1* (0.8 kb) and *AN2* (1kb) were amplified from *Petunia x hybrida* R27 genomic DNA with primers from Table S1 and cloned into pCR-BluntII-TOPO (ThermoFisher). Binding sites were amplified from these plasmids with primers listed in Supplementary Table 1, with the forward primer labelled with Cy5 in 5’. The labelled DNA was purified and incubated with the TnT *in vitro* translation mixture as described in (Silva et al., 2015) before loading on a native acrylamide gel.

### PhDEF protein and antibody production

The *PhDEF* truncated cDNA (without the sequence coding for the MADS domain) was chemically-synthesized with optimization for expression in *E. coli* and cloned into a pT7 expression vector by Proteogenix (www.proteogenix.science). The expected PhDEF protein starts at aminoacid 60 (PSITT…) and ends at the last aminoacid of the sequence (…FALLE), and a 6xHis tag was added at the N-terminal part of the protein. The 6xHis-PhDEF protein was purified by affinity column with a Nickel resin under denaturing conditions (8M urea) by Proteogenix. The purified protein was injected in two rabbits for immunization by Proteogenix, to generate PhDEF-directed polyclonal antibodies, that were purified by affinity against the antigen. Both lots of purified antibodies were validated by Western-Blot in petal or sepal tissues from wt, *phdef-151* and *phtm6* samples.

### Chromatin Immunoprecipitation (ChIP)

Samples (wt: full corolla from 2 flowers; *phdef-151*: second whorl sepals from 3 flowers; *phglo1 phglo2*: second whorl sepals from 3/4 flowers) at stage 8 were collected and ground in liquid nitrogen. Ground tissue was resuspended into 10 mL fixation buffer (10 mM Hepes pH7.6, 0.5 M sucrose, 5 mM KCl, 5 mM MgCl_2_, 5 mM EDTA pH8, Complete Protease Inhibitor Cocktail (Merck), 14 mM 2-mercaptoethanol) and a double cross-linking was performed at room temperature (1 hour with disuccinimidyl glutarate at 2.5 mM with gentle shaking, and 5 minutes with formaldehyde 1%). Cross-linking was stopped by adding glycine at 200 mM and samples were put directly on ice. Cells were lysed with a 40 mL-Dounce tissue grinder (Duran Wheaton Kimble), Triton X-100 was added at 0.6% and the lysate was filtered subsequently through 100 µm and 40 µm nylon meshes to recover nuclei. Nuclei were pelleted for 10 minutes at 3,000 g at 4°C, and the pellet was resuspended in 300 µL of cold nuclear isolation buffer (i.e. fixation buffer without 2-mercaptoethanol), carefully deposited on 600 µL of a 15% Percoll solution (15 % Percoll, 10 mM Hepes pH8, 0.5 M sucrose, 5 mM KCl, 5 mM MgCl_2_, 5 mM EDTA pH8) and centrifuged for 5 minutes at 2,000 g at 4°C. The pellet was resuspended into 900 µL of cold nuclear lysis buffer (50 mM Tris-HCl pH7.5, 0.1% SDS, 10 mM EDTA pH8) to lyse the nuclei, and chromatin was sonicated twice for 15 minutes with a Covaris S220 sonicator (peak power 105, Duty factor 5, Cycles/Burst 200 for 900s). For each sample, 25 µL of magnetic protein-A Dynabeads and 25 µL of magnetic protein-G Dynabeads (Invitrogen) were washed twice with 100 µL of cold ChIP dilution buffer (15 mM Tris-HCl pH7.5, 150 mM NaCl, 1% Triton X-100, 1 mM EDTA pH8) using a magnetic rack (MagRack 6, Cytiva). Beads were mixed with 2.5 µg of anti-PhDEF antibody and 1.8 mL of cold ChIP dilution buffer, and incubated for 2 hours at 4°C on a rotating wheel. Sonicated chromatin was centrifuged for 5 minutes at 15,000 g at 15°C, and 25 µL of supernatant (for wt samples) or 50 µL of supernatant (for *phdef-151* and *phglo1 phglo2* samples) was added to the mix of beads and antibody, and incubated overnight at 4°C on a rotating wheel. Beads were washed twice (one quick wash and one long wash with 15 minutes incubation on a rotating wheel) with each of the following buffers: low salt wash buffer (0.1% SDS, 1% Triton X-100, 2 mM EDTA pH8, 20 mM Tris-HCl pH8, 150 mM NaCl), high salt wash buffer (0.1% SDS, 1% Triton X-100, 2 mM EDTA pH8, 20 mM Tris-HCl pH8, 500 mM NaCl), LiCl wash buffer (0.25 M LiCl, 1% NP40/Igepal,1% deoxycholate, 1 mM EDTA pH8, 20 mM Tris-HCl pH8) and TE buffer. Elution was performed twice with 250 µL of elution buffer (0.1 M NaHCO_3_, 1% SDS) at 65°C. IP and input samples were decrosslinked overnight at 65°C by adding NaCl at 200 mM, then incubating for 2 h at 42°C with 20 µg proteinase K in 10 mM EDTA pH8 and 40 mM Tris-HCl pH6.5. DNA was purified with phenol:chloroform:isoamyl alcohol (25:24:1) followed by chloroform:isoamyl alcohol (24:1), precipitated with ethanol at −20°C and the pellet was washed with ethanol 70 %. The dry pellet was recovered in 50 µL TE and 1 µL was used for each qPCR reaction, which were performed in technical triplicates for each biological replicate (3 for wt and *phdef-151*, 2 for *phglo1 phglo2* and the control without antibody). The qPCR reaction was performed with 1X FastStart Universal SYBR Green (Merck) and 0.3 µM primer mix (Supplementary Table 1), for 40 cycles (15 seconds at 95°C, 1 minute at 60°C) in a QuantStudio 6 Flex instrument (ThermoFisher). Percentage of input was calculated as 100* e(CtIN – log2(DF) – CtIP), with e the efficiency of the primer pair, CtIN the average Ct value for the Input sample, DF the dilution factor and CtIP the average Ct value for the IP sample), as described in (Solomon et al., 2021).

### Accession numbers

Sequence data from this article can be found in the EMBL/GenBank data libraries under accession numbers OQ418981 (AN1), OQ418982 (AN2 and OQ418983 (PhDEF).

## Supplemental Data

**Supplemental Figure 1.** Additional pictures of star and wico flowers.

**Supplemental Figure 2.** Stamens are unfused to the tube in wico flowers.

**Supplemental Figure 3.** Additional pictures of *PhDEF* transcript *in situ* hybridization in wt, star and wico flowers.

**Supplemental Figure 4.** Wt and pink wt flowers observed in the progeny of a star parent.

**Supplemental Figure 5.** Epidermal revertant sectors on star petals.

**Supplemental Figure 6.** Autonomous and non-autonomous effects in star and wico petals. Supplemental Figure 7. Expression of B-class genes and a subset of pigmentation genes in wt, star, wico and *phdef-151* samples.

**Supplemental Table 1.** List of primers used in this study.

**Supplemental Table 2.** Genotyping results of the progeny of a star flower.

**Supplemental Dataset 1.** Differential gene expression calculated by DESeq2.

**Supplemental Dataset 2.** List of the 451 genes downregulated in star and *phdef-151* samples, and not deregulated in wico samples.

## ACKNOWLEDGMENTS

We thank Patrice Bolland, Justin Berger and Alexis Lacroix for plant care assistance, the PLATIM platform (SFR BioSciences Lyon, UAR3444/CNRS, US8/Inserm, ENS de Lyon, UCBL) for electron microscopy technical support, Benjamin Gillet and Sandrine Hugues from the sequencing platform of the Institut de Génomique Fonctionnelle de Lyon for library preparation and sequencing of the transcriptomes of this study, Rémy Belois for assistance for *in situ* hybridization experiments and Daniel Bouyer for assistance for chromatin immunoprecipitation experiments. This work was supported by a PhD fellowship to M.C. from the French Ministry of Higher Education and Research, by a grant to Q.C.S and M.M. from the Agence Nationale de la Recherche (grant ANR-19-CE13-0019, FLOWER LAYER), by a grant to M.M. from IDEXLYON (Université de Lyon, grant ELAN-ERC), and by a grant to V.H. and C.Z. from the Agence Nationale de la Recherche (grant ANR-16-CE92-0023, FLOPINET).

## AUTHOR CONTRIBUTIONS

M.M. and M.V conceived and designed the experiments. M.C., Q.C.S, P.M., P.C., V.H. and S.R.B. performed the experiments. M.C., Q.C.S., J.J., M.V. and M.M. analyzed the data. M.C., C.Z., M.V. and M.M. wrote the article.

