## Supplemental Data for "Cell layer-specific expression of the homeotic MADS-box transcription factor PhDEF contributes to modular petunia petal morphogenesis"

**Supplemental Figure 1.** Additional pictures of star and wico flowers.
**Supplemental Figure 2.** Stamens are unfused to the tube in wico flowers.
**Supplemental Figure 3.** Additional pictures of *PhDEF* transcript *in situ* hybridization in wt, star and wico flowers.
**Supplemental Figure 4.** Wt and pink wt flowers observed in the progeny of a star parent.
**Supplemental Figure 5.** Epidermal revertant sectors on star petals.
**Supplemental Figure 6.** Autonomous and non-autonomous effects in star and wico petals.
**Supplemental Figure 7.** Expression of B-class genes and a subset of pigmentation genes in wt, star, wico and *phdef-151* samples.

**Supplemental Table 1.** List of primers used in this study.
**Supplemental Table 2.** Genotyping results of the progeny of a star flower.

**Supplemental Dataset 1.** Differential gene expression calculated by DESeq2.
**Supplemental Dataset 2.** List of the 451 genes downregulated in star and *phdef-151* samples, and not deregulated in wico samples.

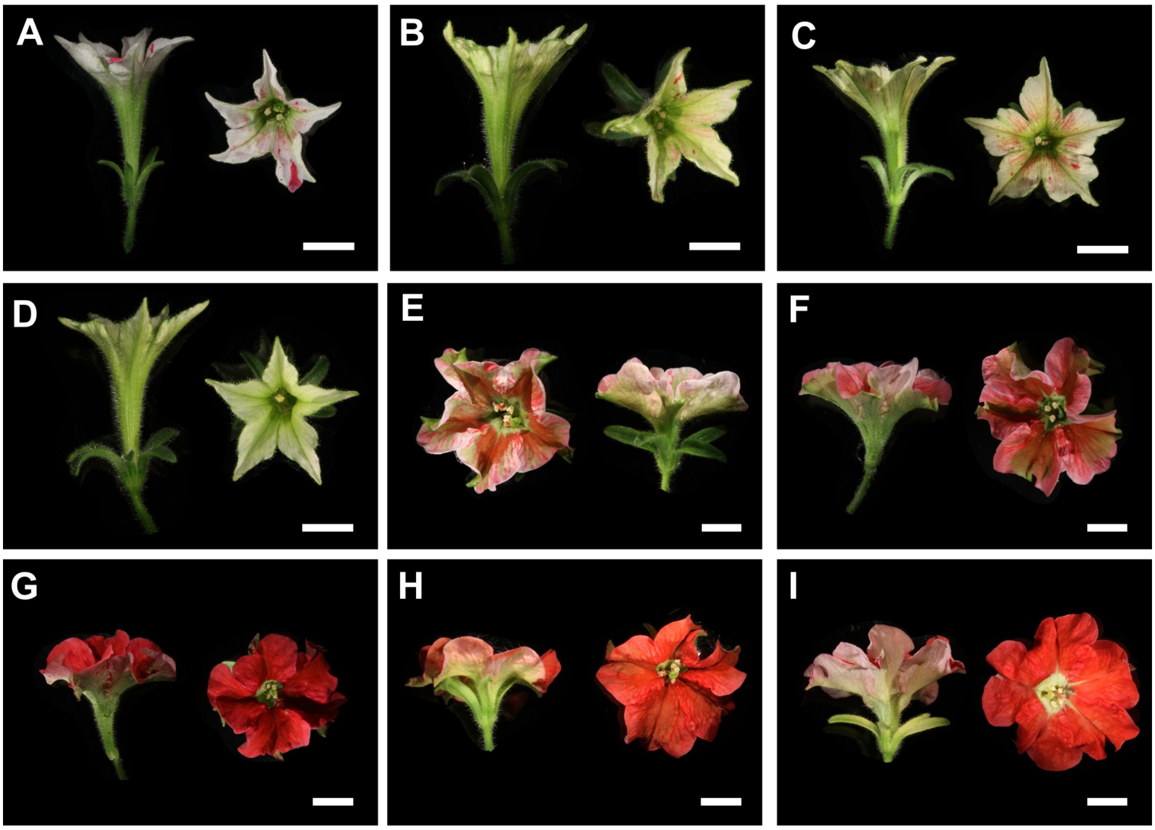


**Supplemental Figure 1.** Additional pictures of star and wico flowers.

Star (A-D) and wico (E-I) flowers from independent branches, viewed from the side (left) and from the top (right). Sepals have been occasionally removed for clarity. Scale bars: 1 cm.

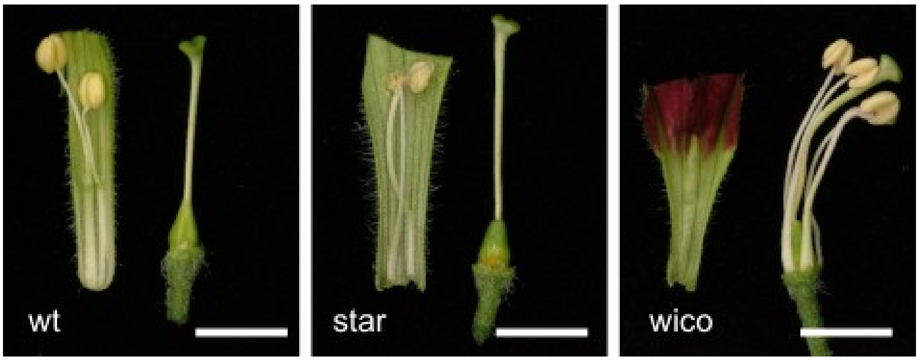


**Supplemental Figure 2.** Stamens are unfused to the tube in wico flowers.

The base of the corolla (left) was detached from the gynoecium (right) in wt, star and wico flowers. In wt and star flowers, the stamens are fused to the petal tube at their base. In wico flowers, the stamens are unfused to the petal tube and therefore remain attached to the flower receptacle. Scale bars: 0.5 cm.

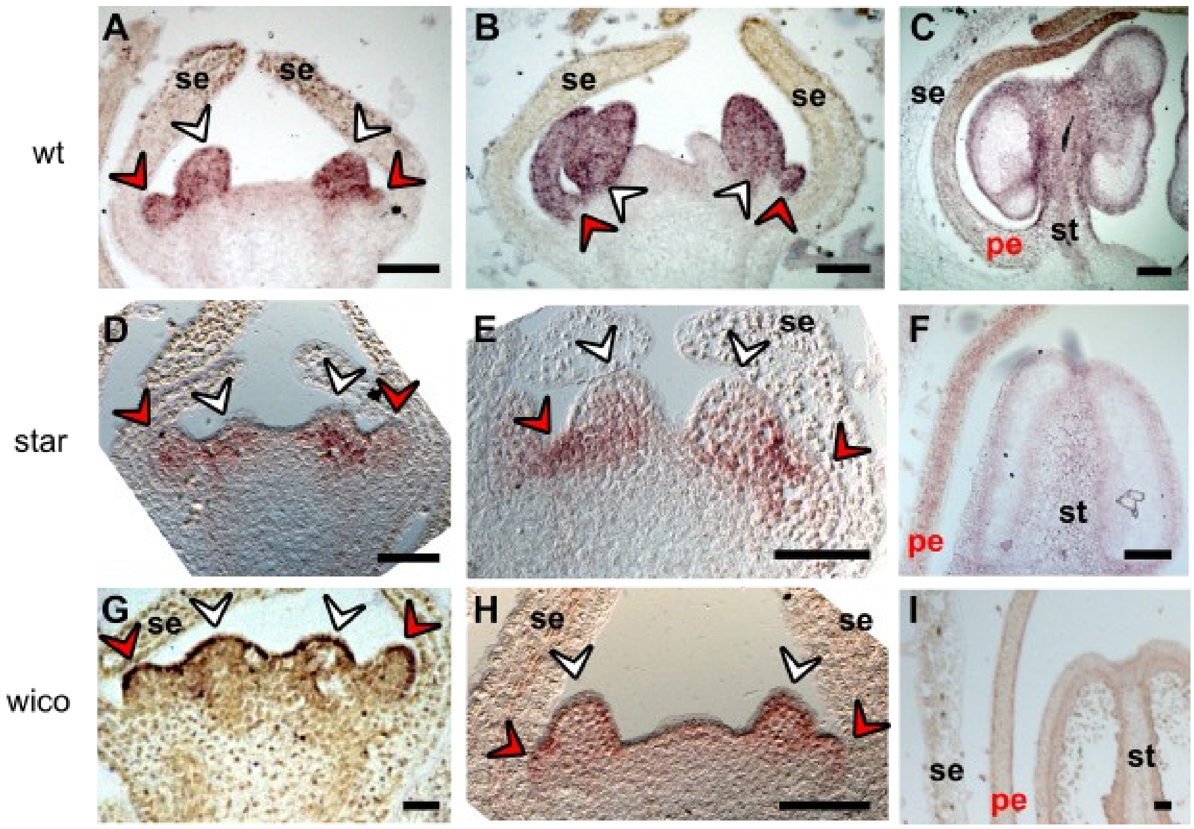


**Supplemental Figure 3.** Additional pictures of *PhDEF* transcript *in situ* hybridization in wt, star and wico flowers.

Longitudinal sections of wt (A-C), star (D-F) and wico (G-I) flowers at various stages (increasing stages from left to right) hybridized with a DIG-labelled *PhDEF* antisense probe. Red and white arrowheads indicate initiating petals and stamens respectively. se: sepals, pe: petals, st: stamen. Scale bar: 20 µm (A, B, D, E, G, H) or 50 µm (C, F, I).

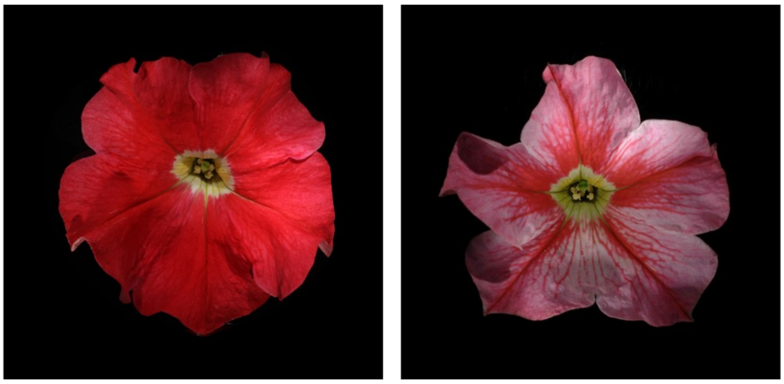

**Supplemental Figure 4.** Wt and pink wt flowers observed in the progeny of a star parent. Pink wt flowers (right) carry a *PhDEF+6* allele and an out-of-frame *phdef* allele, whereas flowers carrying *PhDEF+6* alleles at the homozygous state appear wt (left).

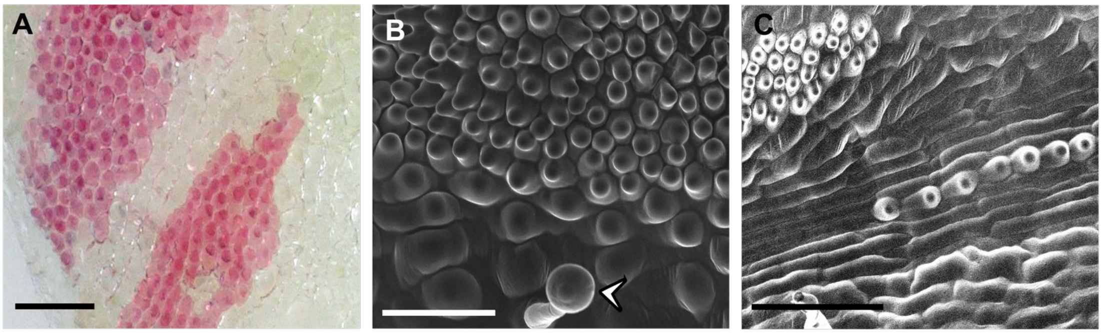


**Supplemental Figure 5.** Epidermal revertant sectors on star petals.

**(A)** Detail of a small pink revertant sector found on a star flower, showing a sharp transition between pigmented and non-pigmented epidermal cells. Scale bar: 100 µm. **(B)** Scanning electron micrograph of the boundary between star and pink revertant petal sectors, showing a quite sharp transition in conical cell shape and size. The white arrow indicates a trichome. Scale bar = 50 µm. **(C)** Scanning electron micrograph of a file of pigmented revertant cells on a star petal limb, located along a vein, showing a sharp transition in conical cell shape. Scale bar: 100 µm.

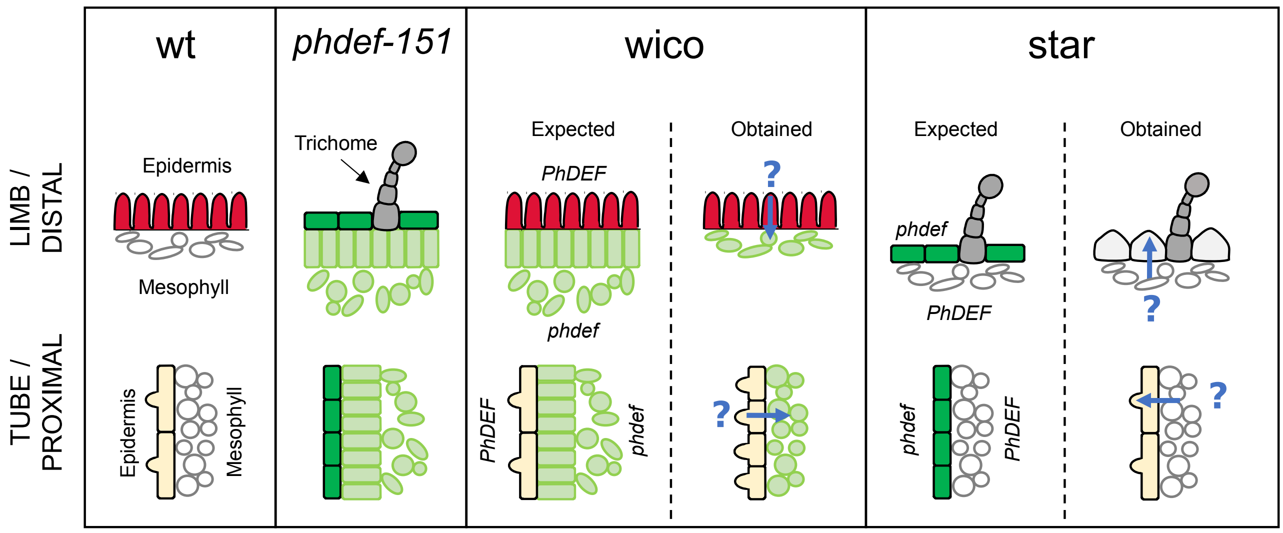


**Supplemental Figure 6.** Autonomous and non-autonomous effects in star and wico petals. Schematic cross-sections of the second whorl organs from wt, *phdef-151*, wico and star flowers, for the distal (limb for petals) and proximal (tube for petals) parts of the organs. Adaxial epidermal cells and mesophyll layer cells are schematized based on cross-sections from Fig. 4, and abaxial epidermal cells have been omitted. For *phdef-151*, the mesophyll is divided in palisade and spongy layers. For wico and star, we propose a schematic view of the expected cross-sections, based on the genotype of the flower, and of the cross-sections that we obtained. The difference between the two suggests the existence of non-autonomous effects (blue arrow) of an unknown nature.

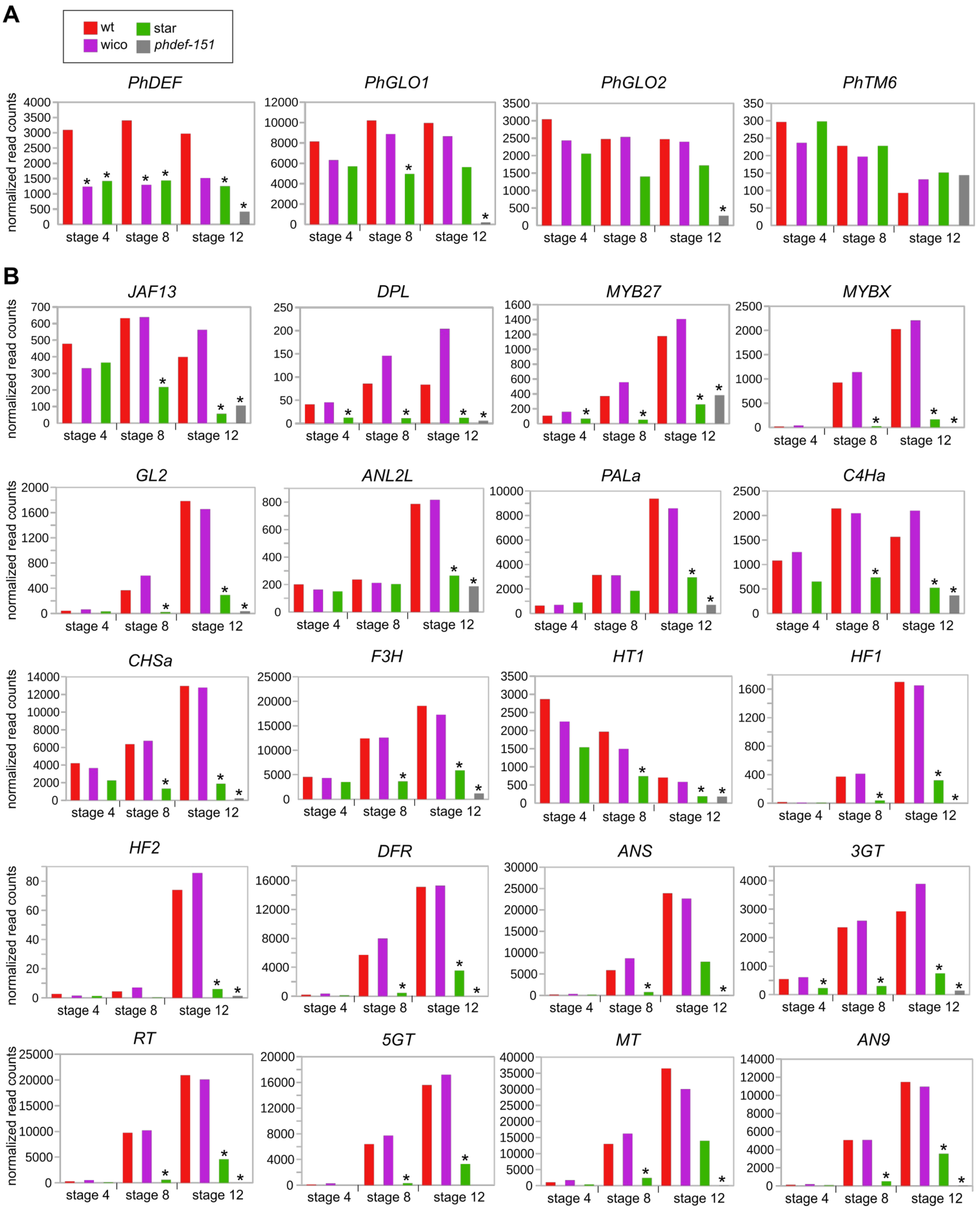


**Supplemental Figure 7.** Expression of B-class genes and a subset of pigmentation genes in wt, star, wico and *phdef-151* samples.

Expression (as normalized read counts calculated by DESeq2) of B-class genes (A) and a subset of pigmentation genes (B) in wt, star, wico and *phdef-151* second whorl organs at stages 4, 8 or 12. The subset of pigmentation genes are the 23 genes significantly down-regulated in star samples (at any stage) and *phdef-151* samples, and not deregulated in wico petals, as in Supplemental Dataset 2. Stars indicate significant down-regulation (log2FC < -1 and adjusted p-value < 0.01).

| Target gene | Application | Primers | 5'-3' sequence |
|---|---|---|---|
| *PhDEF* | Genotyping *phdef-151* | MLY0935<br>MLY0936 | GAACTCACTGTTCTTTGTGATGC<br>GAGAATACTAGTTCTTGGATACGTAC |
| | *In situ* hybridization | MLY1738<br>MLY1739 | GACTAAGCAGTTGTTCGATCTGTAC<br>tgtaatacgactcactatagggcTACTCAAGCAGAGCAAAAGTAGTG |
| | Cloning into pSPUTK | MLY2382<br>MLY2383 | AACATGCCATGGCTCGTGGAAAGATCCAGA<br>AACATGTCTAGACTACTCAAGCAGAGCAAAAGTAGT |
| | ChIP ($PhDEF^{GF1}$) | MLY2626<br>MLY2627 | GCAACACCTTTCACGGTTTTGGCAA<br>CTTGTAGATTTAGAGATAAGATTCTGGAGG |
| *PhGLO1* | Cloning into pSPUTK | MLY2384<br>MLY2385 | AACATGCCATGGGGAGAGGAAAGATAGAGATAAA<br>AACATGTCTAGATTACAACCTCTCCTGCAAATTTGG |
| *AN1* | terminator cloning | MLY2334<br>MLY2335 | CAAGAAGTCAATACACCAGTTAATCCC<br>GCCATCTATTGTTGCATTAGCG |
| | *AN1-bs1* amplification | MLY2336<br>MLY2337 | Cy5-TTTATCCCAAAATAAGTGCCTC<br>AACAAATTTTTGAATTAAAGAGTTAC |
| | *AN1-bs2* amplification | MLY2338<br>MLY2339 | Cy5-TTCACAAACATCAAAGTTTAGC<br>CTAGGGAATAGAGCAAGTAGC |
| | ChIP ($AN1^{GF1}$) | MLY2643<br>MLY2644 | CAAGAATCAATCGTTGTATTAGTCTCTGTCAAAT<br>GCGAATTCCACGAAAAGCGGTGTG |
| | ChIP ($AN1^{GF2}$) | MLY2630<br>MLY2631 | TGTCGACCATTCTTGAACACCTCTCAAAC<br>GGTGCTGGGGCTGGGCCACC |
| | ChIP ($AN1^{GF3}$) | MLY2641<br>MLY2642 | CCTCTGTTCCATAATGAGTGTCTATCTTTTC<br>CCTTTGGTCTAAAGCTAAAAGTGTGTGTAAT |
| *AN2* | terminator cloning | MLY2440<br>MLY2441 | GGGATTTACTTGGTTAATTGGGACC<br>CCAAACATTAAAATCTCTCCAAACTAA |
| | *AN2-bs3* amplification | MLY2450<br>MLY2451 | Cy5-TTATGTACGGTTGTGAGGGA<br>AGGAGAGTTGCGGTAGGC |
| | ChIP ($AN2^{GF1}$) | MLY2737<br>MLY2738 | GGAATGAACTCATAGTTTTAGATGTACCATCG<br>GCCAAACTTGCATGAAAATTACCACAACTAT |
| | ChIP ($AN2^{GF2}$) | MLY2636<br>MLY2637 | CTTCATAAGCTTCTAGGCAACAGGTAAG<br>GAGATGCTAAAGGGGAAGTTCAGGGTT |
| | ChIP ($AN2^{GF3}$) | MLY2634<br>MLY2635 | CCACATAGCAACATATGTAACCCTTTCTTTTT<br>CGATCCCAAATCGAAATAAGTGAGGAG |
| Neg1 (Peaxi162 Scf00207g 00634) | ChIP | MLY2645<br>MLY2646 | GGAGACATAGGATTCACATATCCAACCCAAC<br>GGAGAAATTAAACACTTCCCCACTAGTTGATTA |
| Neg2 (Peaxi162 Scf00164g 00525) | ChIP | MLY2647<br>MLY2648 | CGGCATTCATGGATTTCTTATATTGACCTG<br>TCTAGAAATGACATGAAAAACCATTAGGAGTACT |

**Supplemental Table 1.** List of primers used in this study.

For MLY1739, the T7 Polymerase promoter sequence is in red. For MLY2382 to MLY2385, the XbaI or NcoI restriction sites are in red.

| Plant | Phenotype | Genotype |
|---|---|---|
| 1 | wt | *PhDEF+6 / PhDEF+6* |
| 2 | wt | *PhDEF+6 / PhDEF+6* |
| 3 | wt | *PhDEF+6 / PhDEF+6* |
| 4 | wt | *PhDEF+6 / PhDEF+6* |
| 5 | pink wt | *phdef-151 / PhDEF+6* |
| 6 | pink wt | *phdef-151 / PhDEF+6* |
| 7 | pink wt | *phdef-151 / PhDEF+6* |
| 8 | pink wt | *phdef-151 / PhDEF+6* |
| 9 | pink wt | *phdef-151 / PhDEF+6* |
| 10 | pink wt | *PhDEF+7 / PhDEF+6* |
| 11 | pink wt | *phdef-151 / PhDEF+6* |

**Supplemental Table 2.** Genotyping results of the progeny of a star flower.

11 plants with flowers with a wild-type flower architecture, descendant from a single star flower after selfing, were classified into wt or pink wt phenotypes (see Fig. S3). These plants were genotyped and sequenced at the *PhDEF* locus. All plants with flowers with a wt appearance carried in-frame *PhDEF+6* alleles at the homozygous state. All plants with flowers with a pink wt appearance carried an out-of-frame *PhDEF* allele (*phdef-151* or *PhDEF+7* allele) and an in-frame *PhDEF+6* allele.

**Supplemental Dataset 1.** Differential gene expression calculated by DESeq2.

List of deregulated genes on one-to-one sample comparison with wt. The wt samples are the reference when calculating the fold change (a negative log2FC means that the gene is downregulated in the test sample as compared to wt). BaseMean = mean number of reads across samples, log2FC = log2(fold change), lfcSE = standard error of the log2FC, padj = p-value adjusted for multiple testing, Annotation = functional annotation from the published *P. axillaris* genome, Manual Annotation = manual annotation for MADS-box genes (highlighted in green) and anthocyanin-related genes (highlighted in pink). Genes are considered to be deregulated if log2FoldChange > 1 or < -1, and p-adjusted value < 0.01. The list of all manually annotated MADS-box genes and anthocyanin-related genes (see Methods) can be found in the last spreadsheet.

**Supplemental Dataset 2.** List of the 451 genes downregulated in star and *phdef-151* samples, and not deregulated in wico samples.

The wt samples are the reference when calculating the fold change (a negative log2FC means that the gene is downregulated in the test sample as compared to wt). log2FC = log2(fold change), only reported here if > 1 or < -1 and p-adjusted value < 0.01; Annotation = functional annotation from the published *P. axillaris* genome; Manual Annotation = manual annotation for anthocyanin-related genes (highlighted in pink) and MADS-box genes. Genes are considered to be downregulated if log2FoldChange < -1, and p-adjusted value < 0.01. Only genes downregulated in star samples at least at one stage, also downregulated in *phdef-151* samples, and not deregulated in wico samples at any stage, are shown here.
